# Landscape of human protein-coding somatic mutations across tissues and individuals

**DOI:** 10.1101/2025.01.07.631808

**Authors:** Huixin Xu, Rob Bierman, Dayna Akey, Cooper Koers, Troy Comi, Claire McWhite, Joshua M. Akey

## Abstract

Although somatic mutations are fundamentally important to human biology, disease, and aging, many outstanding questions remain about their rates, spectrum, and determinants in apparently healthy tissues. Here, we performed high-coverage exome sequencing on 265 samples from 14 GTEx donors sampled for a median of 17.5 tissues per donor (spanning 46 total tissues). Using a novel probabilistic method tailored to the unique structure of our data, we identified 8,470 somatic variants. We leverage our compendium of somatic mutations to quantify the burden of deleterious somatic variants among tissues and individuals, identify molecular features such as chromatin accessibility that exhibit significantly elevated somatic mutation rates, provide novel biological insights into mutational mechanisms, and infer developmental trajectories based on patterns of multi-tissue somatic mosaicism. Our data provides a high-resolution portrait of somatic mutations across genes, tissues, and individuals.

## Introduction

Somatic mosaicism, a phenomenon where different cells within the same individual have different genetic sequences, plays pivotal roles in human biology. This stands in contrast to germline variants, which are inherited from parents to offspring and are therefore present in all cells. Such mutations are intimately associated with aging and cancer, and over 30 human diseases have been identified as resulting from somatic mutations (1–4).

Furthermore, they provide valuable insights into human development and the complex processes that shape our bodies and health (5–7).

Somatic mutations are pervasive and can occur at every stage of human development and throughout life. A human zygote undergoes myriad cell divisions to result in a fully differentiated multicellular organism, creating numerous opportunities for somatic mutations to occur. Even terminally differentiated cells continue to acquire somatic variants in their genomes through non-replicative processes. As estimated by Lynch (8), the rate of somatic mutations is 4-25 times higher than that of germline mutations, leading to a theoretically enormous burden of somatic mutations in humans.

Our current understanding of somatic mosaicism is largely derived from the study of cancer genomics (9). However, it remains unclear whether the somatic mutations observed in cancer are representative of those in healthy individuals. Further investigations into the mechanisms underlying mutations in normal tissues could reveal new opportunities for early detection of diseases and the establishment of reference lines distinguishing pathological from benign somatic variations.

Recent studies have expanded our knowledge of somatic mutations in various normal tissues, including skin (10–12), esophagus (13,14), brain (15–17), blood (18), endometrial glands (19–21), colon (22,23), liver (24,25), breast (26), bronchial epithelium (27) and stomach (28). These studies have uncovered a large number of mutations in normal tissues and have revealed clonal expansions in healthy tissues, heterogeneity in the rates of mutations within the genomes of individuals, mutational signatures, and positive selection of cancer genes. In addition to providing insights into mutational processes and levels of somatic mosaicism across tissue and individuals, these findings could also have significant implications for the understanding of the pathobiology of cancer, other diseases, and therapeutic interventions. However, these studies have been limited to one or a few tissues per individual.

The two main impediments to a more comprehensive understanding of human somatic mosaicism are the lack of appropriate data sets where multiple tissues from various individuals have been sampled and powerful statistical methods tailored to the unique structure of this data. The Genotype-Tissue Expression (GTEx) project offers a valuable resource for investigating somatic mutations in normal tissues (29). It has provided RNA-seq data from varied tissues collected from a common set of donors, which has been leveraged to identify somatic mutations (30–32). Despite the insights gained from previous studies based on GTEx RNA-seq, challenges and limitations in somatic mutation detection in RNA-seq persist, leaving important questions unanswered in the exploration of somatic variation patterns (33).

Although several studies have used the GTEx data to study somatic mutations, they all identified somatic mutations in RNA-Seq data, which is potentially error-prone (30–32). To this end, we performed high-coverage exome sequencing in 265 GTEx samples, spanning 46 distinct tissues collected from 14 individuals. We also developed a novel method, referred to as multi-tissue SOmatic Mutation Analyzer (mSOMA), for detecting somatic mutations in multiple normal tissues from the same individuals. In total, we identified 8,470 somatic mutations across all 265 tissue samples and leveraged the compendium of somatic mutations to better understand their patterns, characteristics, and determinants. Our data provides important novel insights into the burden of somatic mutations across tissues and individuals.

## Results

### Overview of study

We studied somatic mutations in 14 individuals, referred to as donors hereafter, that were part of the GTEx Project (Fig. 1A) (29). The GTEx Project collected DNA, RNA, and other molecular phenotypes in multiple tissues from each donor as well as extensive covariate data, providing a powerful resource to better understand the patterns and determinants of somatic variability (Fig. 1A). For this purpose, we performed high-coverage exome sequencing on a total of 265 samples, spanning 46 tissues across all 14 donors (Fig. 1B), with an average of 18.9 tissues sequenced per donor (Fig. 1B and 1C; Table S1). The tissues sequenced span a broad range of body systems and all three germ layers (Fig. 1B). Of the 14 donors, eight are female and six are male (Fig. 1C & Table S1). Donor age ranges from 20 to 69 years old, with a mean of 51.7 years old (Fig. 1C), and 50% (7/14) of donors are over 60 years old (Fig. 1C & Table S1). The mean depth of captured exome regions before and after quality control is 175x and 140x (Fig. S1A), respectively. On average, the cumulative coverage across all tissues sequenced per donor was 2,649x (ranging from 1,454 to 4,260x; Fig. 1D & Table S1).

**Figure 1.**
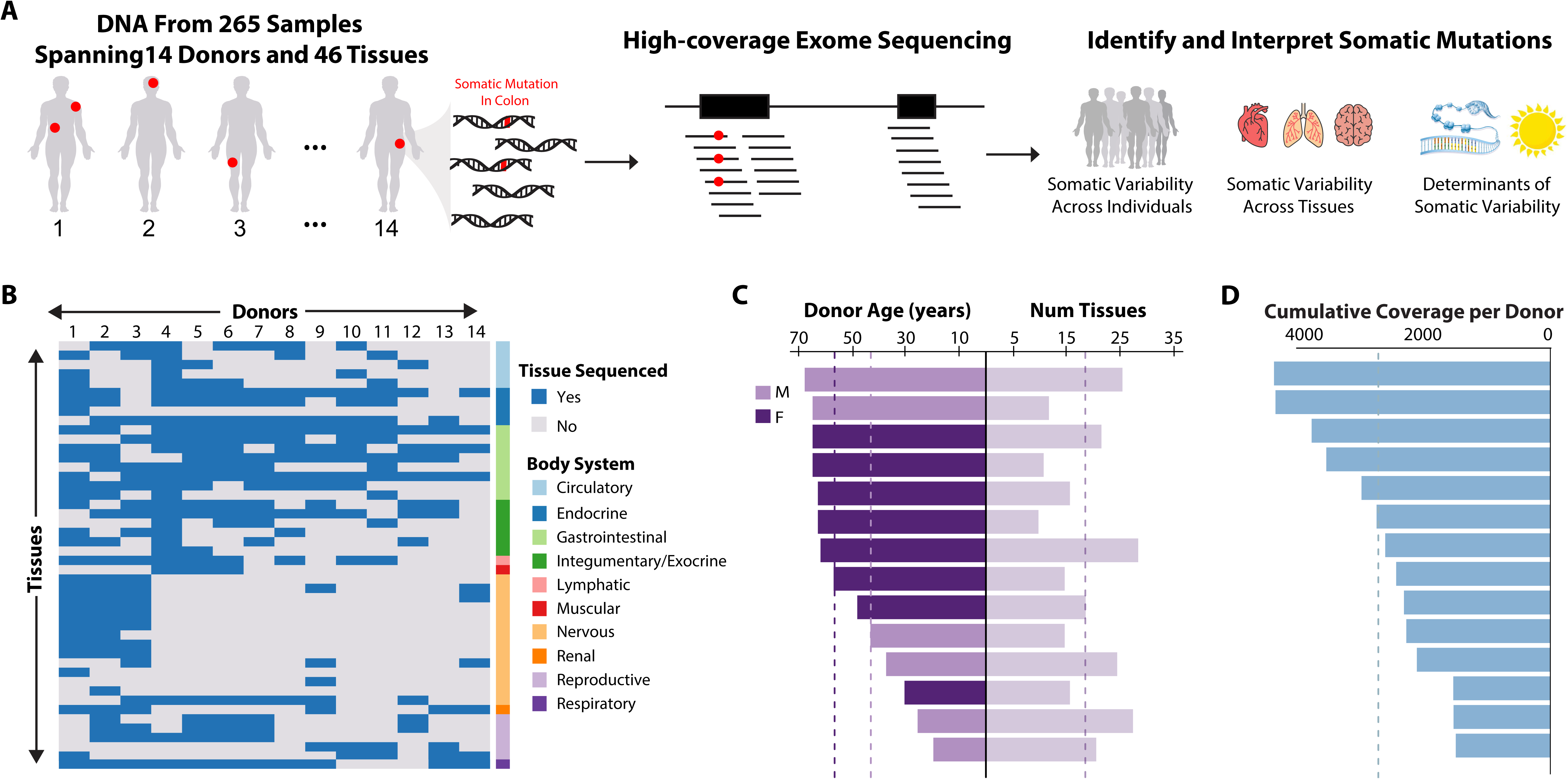
Overview of study design to identify somatic mutations from 265 high- coverage exome sequences. **(A)** Schematic illustration of study design to identify somatic mutations (red circles) in 14 GTEx donor individuals. High-coverage exome sequencing was performed on DNA isolated from a median of 17.5 tissues per donor (in total, 265 samples were analyzed, spanning 46 distinct tissues across all donors). The set of identified somatic mutations was leveraged to test hypotheses about the burden and determinants of somatic mosaicism across individuals and tissues. **(B)** The heatmap summarizes the tissues (rows) analyzed in the 14 donors (columns). Tissues are organized into body systems, as indicated on the right side of the heatmap. Blue and gray-filled rectangles indicate whether DNA from a particular tissue was or was not sequenced, respectively, in a particular donor. **(C)** Barplots indicate the age and sex (left) and the number of tissues sequenced per donor (right). Dashed lines indicate the mean age for males and females (left) and the mean number of tissues sequenced per donor (right). **(D)** The cumulative amount of sequence coverage across all tissues studied in each donor. The dashed line shows the average coverage of 2,800X across donors.

### A novel method to identify somatic mosaicism in multi-tissue study designs

Most existing methods to detect somatic single nucleotide variants (SNVs) were designed in the context of cancer genomics (34–38). These methods typically require a test tumor sample and a matched reference sample from the same individuals. In such analyses, somatic mutation calls are discerned by sequencing both tumor and non-cancerous tissue pairs from the same individual and then filtering calls that are common to both test and reference tissues, as these are likely to be germline. However, these methods are not suitable for our data, which comprises various tissues sequenced per individual without a distinct reference sample.

Therefore, we developed mSOMA, a reference-free somatic SNV calling method that is specifically tailored to the unique structure of our data (Fig. 2A; STAR Methods). Briefly, after applying read mapping, standard GATK best practice (36) and pre-filters, we modeled alternative allele counts by Beta-binomial distributions. Here, 96 combinations of trinucleotide contexts were considered independently to capture sequence-composition-specific error rates. Somatic variants that are potential true positives (i.e., those with a higher number of reads supporting a variant allele than a local background error rate) will be identified against this background distribution using hypothesis testing. For instance, with reference and alternative alleles denoted as C and A respectively at a genomic location (Fig. 2A), the null hypothesis posits that the additional presence of alternative allele A is attributable to sequencing errors or other technical artifacts. The alternative hypothesis proposes that the occurrences of additional alternative allele A are due to somatic variants.

**Figure 2.**
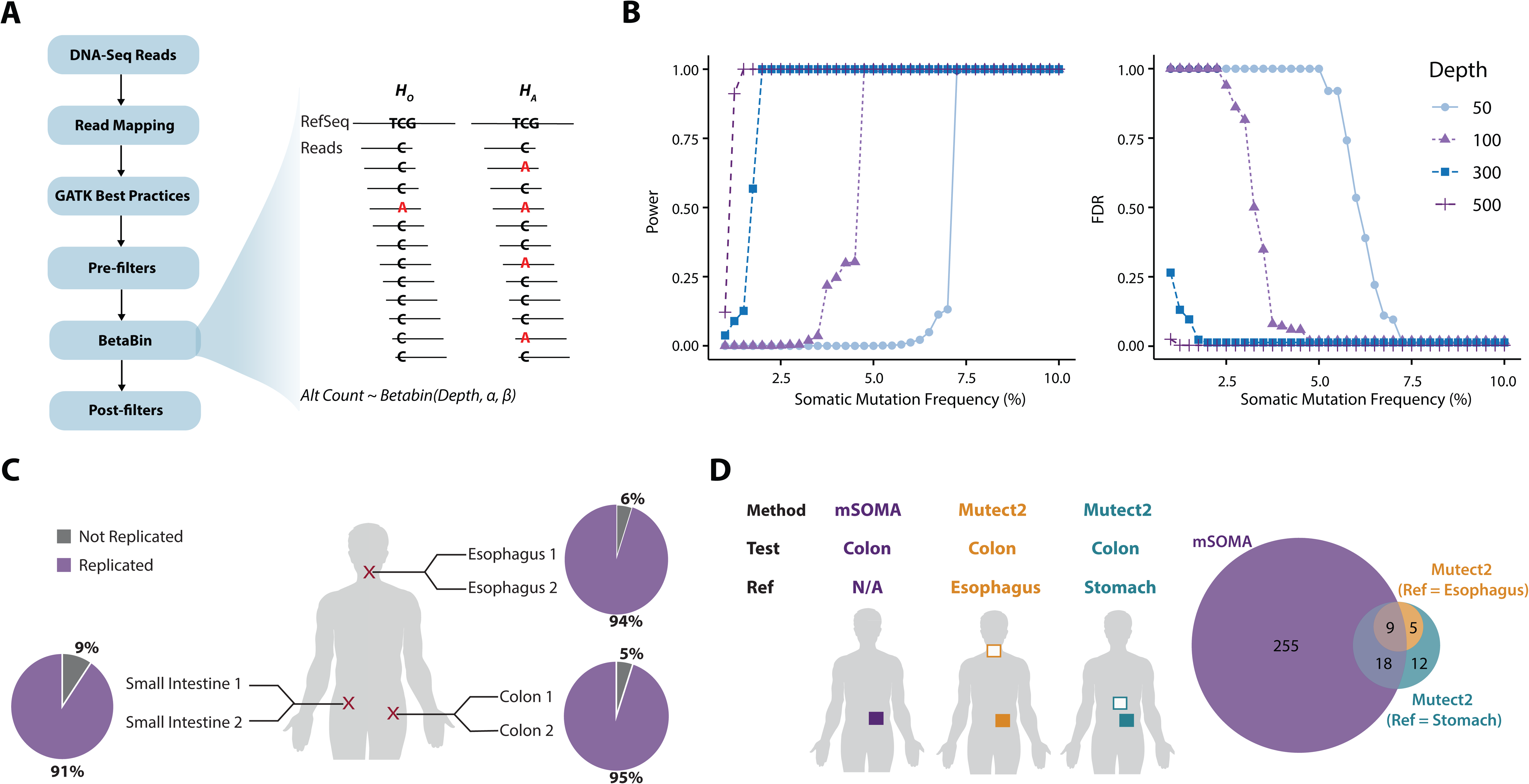
SOMA is a powerful approach to identify somatic mutations in multi-tissue study designs. **(A)** Summary of how SOMA identifies somatic SNVs. We first perform read mapping, identify (and subsequently mask) germline variants with GATK, and apply a series of pre-filters. Next, we model the number of alternative allele counts (*AC*) at a locus under the null hypothesis as a beta-binomial random variable. To account for error rates that vary across different sequence contexts, we inferred 96 distinct null beta-binomial distributions (corresponding to the six distinct mutational types and the immediate flanking base up and downstream of the mutation). We then formally test whether there are more alternative alleles than expected under the null hypothesis, given the coverage, mutational type, and sequence context. Additional post-filters are applied to obtain a final, refined call set. **(B)** The power (left) and FDR (right) of SOMA as a function of somatic mutation frequency and sequencing depth. **(C)** Replication of calls in three pairs of duplicate samples. One random sample from each duplicate pair was used to discover somatic mutations and the other sample was used for validation. Pie charts indicate the percentage of somatic mutation calls that were (purple) or were not (gray) replicated for each pair of duplicate samples. **(D)** Comparison of somatic mutation calls made by SOMA (purple) and Mutect2 (orange and green) in ten transverse colon samples. Mutect2 was performed using the esophagus (orange) or stomach (green) from the same donor as reference samples.

The test statistic is the observed Alternative allele Count (*AC*). Under the null hypothesis, *AC* follows a Beta-binomial distribution, where α and β were inferred independently for each sequence context. *P*-value will be calculated independently for each locus. Finally, additional post-filters are applied to yield a reliable and refined call set (STAR Methods).

### Somatic variants are accurately detected using mSOMA in bulk tissue DNA-seq data

We performed simulations based on allele counts derived from real data to study the relationship between somatic SNV frequency and both power and false discovery rate (FDR) at different sequencing depths (STAR Methods). The simulation was performed by varying the somatic mutation frequency and calculating the power and FDR at different sequencing depths (50x, 100x, 300x and 500x). As expected, the power and FDR were positively and negatively correlated, respectively, with somatic mutation frequency and sequencing depth (Fig. 2B). For example, when the somatic SNV frequency stands at 2%, the power exhibits a range of 0%, 12.1% and 100% at sequencing depths of 100x, 200x and 300x, respectively.

When the somatic variant frequency increases to 7.5%, power reaches 100% for all sequencing depths. Conversely, our analysis of mSOMA demonstrates a consistent decrease in FDR with elevated somatic mutation frequency and sequencing depth (Fig. 2B). For situations where the somatic variant frequency is equal to or exceeds 2%, coupled with a coverage of 200x or higher, the FDR remains at or below 18.9%. FDR remains zero for depths of >=300x but increases sharply to nearly 100% at lower depths. This underscores mSOMA’s heightened ability to accurately identify common somatic variants using bulk DNA sequencing as both the mutation frequency and sequencing depth increase. It also indicates mSOMA’s capacity to maintain a low rate of false positives under these specific conditions.

In our study, we sequenced three pairs of technical replicate tissue samples using identical DNA libraries to examine the reproducibility of somatic variations calls (STAR Methods). One duplicate from the pairs was randomly selected for somatic variant discovery using mSOMA, while the second served as the validation sample to verify the identified somatic SNVs. To assess the concordance between discovery and validation samples, we calculated the replication rate, which is the percentage of mutation calls observed in the discovery sample that had at least one high-quality read supporting the variant allele in the validation sample. The overall replication rate using mSOMA was found to be 93% (140/150), exhibiting a range from 91% to 95% (Fig. 2C). To further refine our results, we replaced three discovery samples in a cohort of 265 samples with validation samples, subjecting them to additional post-filters to obtain refined calls for subsequent comparisons.

We observed that variants with a higher weighted variant allele frequency (VAF) shared an increased Jaccard similarity between duplicates (Fig. S1B). Specifically, the Jaccard similarity increased from 0.205 at a weighted VAF of 2%, to 0.5 when the weighted VAF reached 7.5%. This indicates that our method is effective and consistent in identifying true somatic variants, especially at higher frequencies.

Finally, to compare performance of different somatic SNV callers, we applied mSOMA to analyze 10 colon samples and compared the outcomes with Mutect2 (39) (STAR Methods). Given the absence of a clear test-reference structure in our data, with no evident reference sample available, we investigated the impact of reference sample selection on the identification of somatic mutations using Mutect2. Specifically, we treated esophagus and stomach tissues from the same donors as matched-normal references. Notably, mSOMA demonstrated a substantial increase, detecting 6-19 times more candidate SNVs compared to Mutect2 (Fig. 2D). Despite a modest overlap, mSOMA was capable of detecting 60-64% of the calls made by Mutect2 (Fig. 2D), underscoring its sensitivity in identifying somatic mosaicism. It is crucial to highlight that Mutect2 exhibited variability in call sets when different reference samples were utilized. This emphasizes the limitation of Mutect2 and other reference-based somatic SNV calling approaches in our study, where multiple tissues from the same donors were sequenced, and a distinct reference tissue sample was not available.

### A compendium of somatic mutations and their characteristics

We identified 8,470 autosomal somatic SNVs across 265 samples using mSOMA (Fig. 3A & Fig. S1C), spanning 46 tissues and 14 donors. The median VAF was found to be 2.3%, with a range from 0.6% to 17.2% (Fig. 3B). As expected, there was a negative correlation between DNA sequencing coverage and VAF (Spearman *R* = -0.91; Fig. S1D). This correlation arises from the increased likelihood of identifying variants with low allele frequency in sites with high coverage. Remarkably, the presence of somatic mosaicism was ubiquitous, with 100% of donors and 100% of tissue samples displaying detectable levels of somatic SNVs.

**Figure 3.**
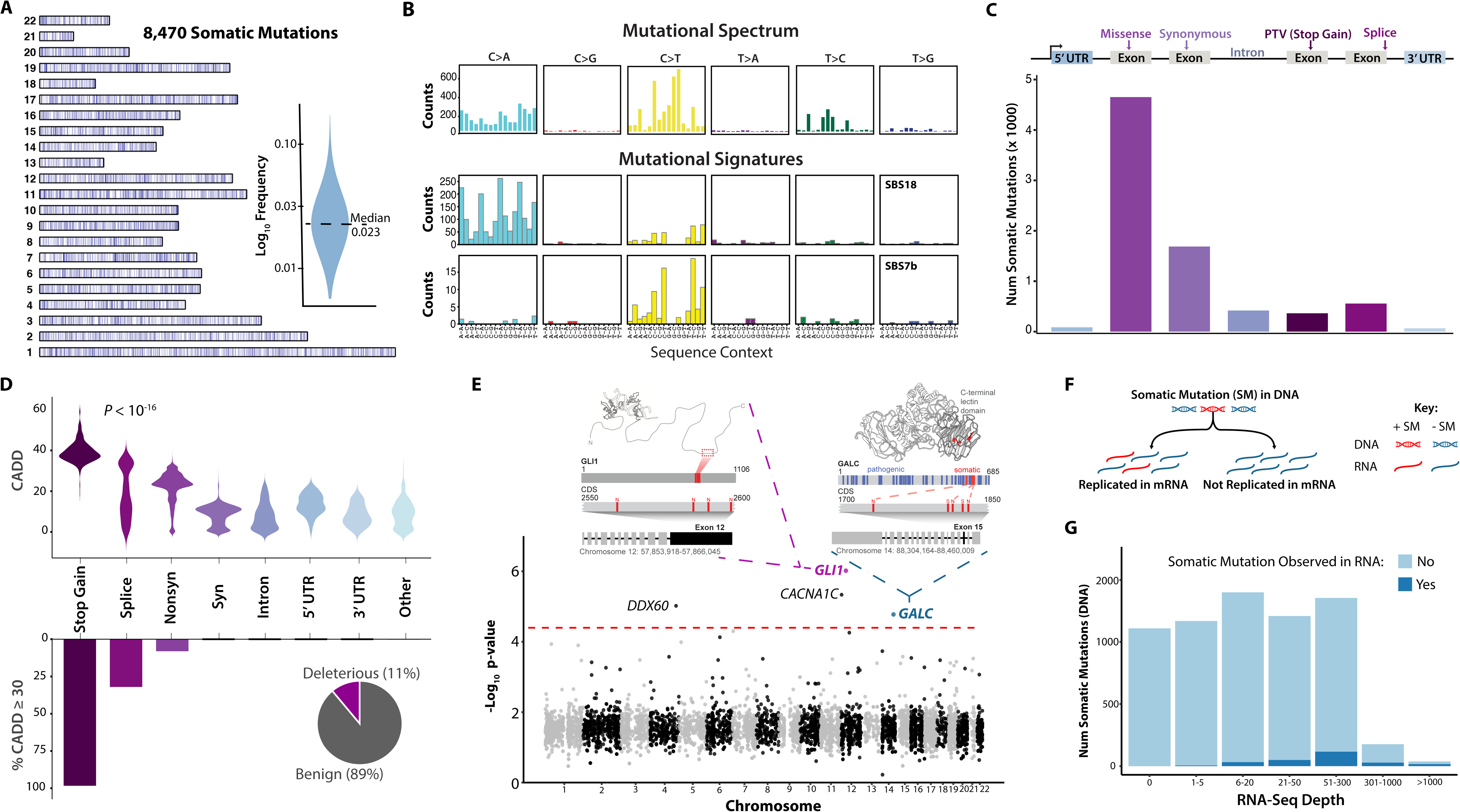
A compendium of somatic mutations and their characteristics. **(A)** Genomic distribution of the 8,470 identified somatic mutations. The inset shows the distribution of somatic mutation frequency (log_10_ transformed). **(B)** Summary of the observed mutational spectrum and inferred mutational signatures. The mutational signature SBS18 is identified in all tissues, whereas SBS7b is specific to sun-exposed skin. **(C)** The number of somatic mutations across functional classes, with gene structure elements depicted at the top. **(D)** Identifying putatively deleterious somatic mutations. The upper panel shows that the distribution of CADD scores significantly varies across functional classes (*P*-value obtained from a Kruskal–Wallis test). The lower panel illustrates the proportion of deleterious somatic mutations (defined as those with a CADD score ≥30) across functional classes. The pie chart summarizes the overall proportion of putatively deleterious and benign somatic mutations. (**E**) Identifying clusters of somatic mutation across the exome. The x-axis of the Manhattan plot represents position in the exome and the y-axis corresponds to the -log_10_ *P* value derived from a Poisson test for enrichment of somatic mutations in non-overlapping 50bp windows. The dashed red dashed line indicates an FDR threshold of 10%. The insets illustrate examples of somatic mutation hotspots in exon 12 of *GLI1* and exon 15 of *GALC*. The gene structure, somatic mutations in the coding sequence (N and S denote nonsynonymous and synonymous mutations, respectively), and pathogenic variants (blue) are shown for each example. **(F)** Schematic illustration of testing whether somatic mutations identified in DNA (red) are observed in mRNA from the same tissue (red and black denote transcripts that do and do not carry the somatic variant). **(G)** Observed distribution of DNA identified somatic mutations that are observed in RNA-Seq.

### Mutational signatures across normal tissues

Our analysis of the mutational spectrum showed that C>T (46.2%) and C>A (33.7%) SNVs were the most prevalent, followed by T>C (13.8%) mutations (Fig. 3C). The least frequent mutational type was C>G (1.2%). Using SignatureAnalyzer (40) to extract mutational signatures (STAR Methods), we found that SBS18, which is primarily characterized by C>A variants, occurs in all tissues as a whole (Fig. 3C & Table S2). This signature could be indicative of DNA damage caused by reactive oxygen species. SBS18 has been previously detected in various cell types of normal tissues (41). We also identified SBS7b in sun-exposed skin, which is likely due to exposure to ultraviolet light. SBS7b is marked by a predominance of C>T transitions (Fig. 3C), particularly when the mutated cytosine is preceded by another pyrimidine (in a TpC or CpC context).

Additionally, SignatureAnalyzer (40) also unveiled the widespread occurrence of SBS15 or SBS6 across normal tissues (Table S2). While the unique nature of our calls - mostly common somatic variants based on bulk DNA - may reveal new mutational signatures compared to other studies on normal tissues, we suspect that SBS15 or SBS6 might be confounded by SBS1, as reported by (42). This suspicion arises from the fact that SBS15 or SBS6 is associated with defective DNA mismatch repair and microsatellite instability (MSI), and is characterized by N[C[>[T]G. Consequently, we propose that SBS15 or SBS6 and SBS1 may not be distinguishable, consistent with a previous study (42), and the reported SBS15 or SBS6 represents a mixture of SBS1 and other contributing factors.

### Unexpectedly high ratio of missense to synonymous somatic mutations

Somatic mutations exhibit diverse functional classes. Specifically, 4.4% (376/8470) of these mutations result in stop-gained alterations, while 2.4% (200/8470) are splice variants (Fig. 3D). 13.5% (1146/8470) of identified somatic SNVs occur within intronic regions (Fig. 3D). Among the identified mutations, a substantial proportion - 56.4% (4774/8470) - are missense mutations, while synonymous variants account for 20.7% (1750/8470) (Fig. 3D).

Thus, the observed ratio of missense to synonymous SNVs is 2.73 (4774/1750). To study the distributions of the expected ratio of missense to synonymous somatic mutations, we investigated various mutational models, such as generating random mutations in genes under the assumption that each mutation is equally likely, or adhering to patterns of observed somatic variations within specific contexts (STAR Methods). Surprisingly, none of these mutational models fully captured the patterns evident in the empirical ratio of missense to synonymous somatic mutations (Fig. S1E).

### Characteristics of deleterious somatic variants

We employed Combined Annotation Dependent Depletion (CADD) (43) to assess the functional impacts of somatic mutations. CADD is a measure that integrates various genomic features to assign a Phred-scaled score to each variant, predicting its deleteriousness.

Significant differences were observed in the distributions of CADD scores across various functional classes (*P* < 10^-10^, Kruskal–Wallis Test; Fig. 3E top). Somatic mutations were classified as deleterious if their CADD score was 30 or higher. In general, 11% of somatic SNVs were found to be deleterious (Fig. 3E). There was no significant difference in VAFs between deleterious and benign variants (*P* = 0.77, Wilcoxon Test; Fig. S1F), suggesting a lack of widespread evidence for purifying selection, consistent with previous findings (41,44).

A substantial portion of stop-gained (98.4%) and splice (94.0%) somatic variants are predicted to have deleterious effects (Fig. 3E bottom). Among missense somatic SNVs, 8.3% were deemed deleterious. In contrast, no deleterious effects were associated with synonymous, intron, or Untranslated Region (UTR) somatic mutations (Fig. 3E bottom).

### Identifications of somatic variant clusters

We further explored the local distribution of somatic variants within exomes. In pursuit of this, we segmented the coding sequence (CDS) of each gene into nonoverlapping 50-base-pair windows and investigated whether any of these windows exhibited an enrichment of somatic variants (STAR Methods). Our observations revealed a non-uniform distribution of somatic variants across the exome, with notable clusters emerging in specific genomic regions (Fig. 3F). Remarkably, our investigation pinpointed four distinct 50-bp genomic regions in the genes *GLI1*, *CACNA1C*, *DDX60*, and *GALC* that exhibited significant somatic mutation clusters (*P* = 5.2×10^-7^, 4.6×10^-6^, 9.5×10^-6^ and 1.7×10^-5^, respectively; Poisson Test; Fig. 3F).

We detected four missense mutations in the *GLI1* gene in sun-exposed skin, non-sun-exposed skin, prostate, and testis from four unrelated individuals. Interestin3gly, all these four SNVs were concentrated between the 2550th and 2600th positions of the CDS of *GLI1* (Fig. 3F), specifically within the 12th exon. *GLI1*, recognized as an oncogene, encodes a zinc finger protein (45). This transcription factor, activated by the sonic hedgehog signal transduction cascade, regulates stem cell proliferation. The activity and nuclear localization of this protein are negatively regulated by p53 in an inhibitory loop. Given its pivotal role, *GLI1* serves as a promising therapeutic target for cancer (45). Another cluster example involves four somatic variants, including two pathogenic missense and two synonymous mutations, that are concentrated between the 1800th and 1850th positions of the CDS of *GALC* (Fig. 3F). These insights into localized somatic variant distributions shed light on potential functional implications and highlight specific genomic regions of interest for further investigation.

### Limitations of identifying somatic mutations from RNA-Seq

We undertook an analysis to assess the agreement of somatic mutation calls between DNA-seq and RNA-seq (STAR Methods). Of the cohort studied, 80.7% (214/265) of the DNA-seq samples, in which 7,059 somatic SNVs were identified, had corresponding RNA-seq samples. The validation criterion for a somatic variant found in DNA-seq was met if at least one identical variant allele was present in the matched RNA-seq sample. The validation rate was determined as the percentage of DNA somatic variants that were confirmed using RNA-seq.

Notably, 24.4% (1,721/7,059) of somatic SNVs were not expressed in the RNA-seq samples (Fig. 3G). Our exploration further revealed that the validation rate of somatic mutations identified in DNA increased in correlation with the relative transcript abundance in the matched RNA-seq sample (Fig. 3G). Specifically, only 1.0% (13/1,304) of somatic mutations found in the DNA-seq were observed in the RNA-seq data when its coverage was below 5x (Fig. 3G). As the RNA coverage increased, validation rates saw a notable surge, reaching 18.4% for somatic variants discovered in DNA when the coverage ranged between 50x and 300x. The most substantial validation rates, escalating to 67.1% (47/70), were achieved when the RNA coverage exceeded 1,000x (Fig. 3G). Moreover, our analysis showed a positive correlation between the validation rate of somatic SNVs identified in DNA-seq and their VAFs (Fig. S1G). When the RNA-seq data exhibits substantial coverage, exceeding 1000x, and the DNA-seq data demonstrates a VAF greater than 3%, the validation rate for somatic variants detected in DNA-seq can reach an impressive 87% (13/15). It is important to note that the extracted DNA and RNA come from different cells and thus spatial heterogeneity in the location of a somatic mutation in a tissue may contribute to differences observed in the RNA and DNA data.

### Somatic mutations shared across multiple tissues exhibit distinct characteristics compared to unique somatic mutations

In our study, we identified 95 somatic mutations that were shared across more than one tissue within a single donor (STAR Methods). For instance, a stop-gained C>T somatic SNV in *ASXL1* was observed across three tissues: the Minor Salivary Gland, Spleen, and Adrenal Gland (Fig. 4A), from a donor of 64 years old. This mutation truncates the wild-type *ASXL1* protein of 1541 amino acids (wild type C) into a shorter one of 417 amino acids (mutated type T). We further investigated whether these shared somatic mutations exhibited similar characteristics to unique somatic variants, which are variants observed in only one tissue. Our findings revealed that shared somatic variants had slightly but significantly higher allele frequencies compared to unique somatic SNVs (3.0% vs. 2.5%; *P* = 7.4×10^-4^, Wilcoxon Test; Fig. 4B & Fig. S1D). Additionally, shared somatic mutations displayed marginally lower CADD scores (15.9% vs. 18.3%; *P* = 4.4×10^-2^, Wilcoxon Test; Fig. 4C). This suggests that somatic mosaicism that occurs earlier in development is less constrained and not subject to selection pressure, a finding that aligns with the results from Rockweiler et al. (32).

**Figure 4.**
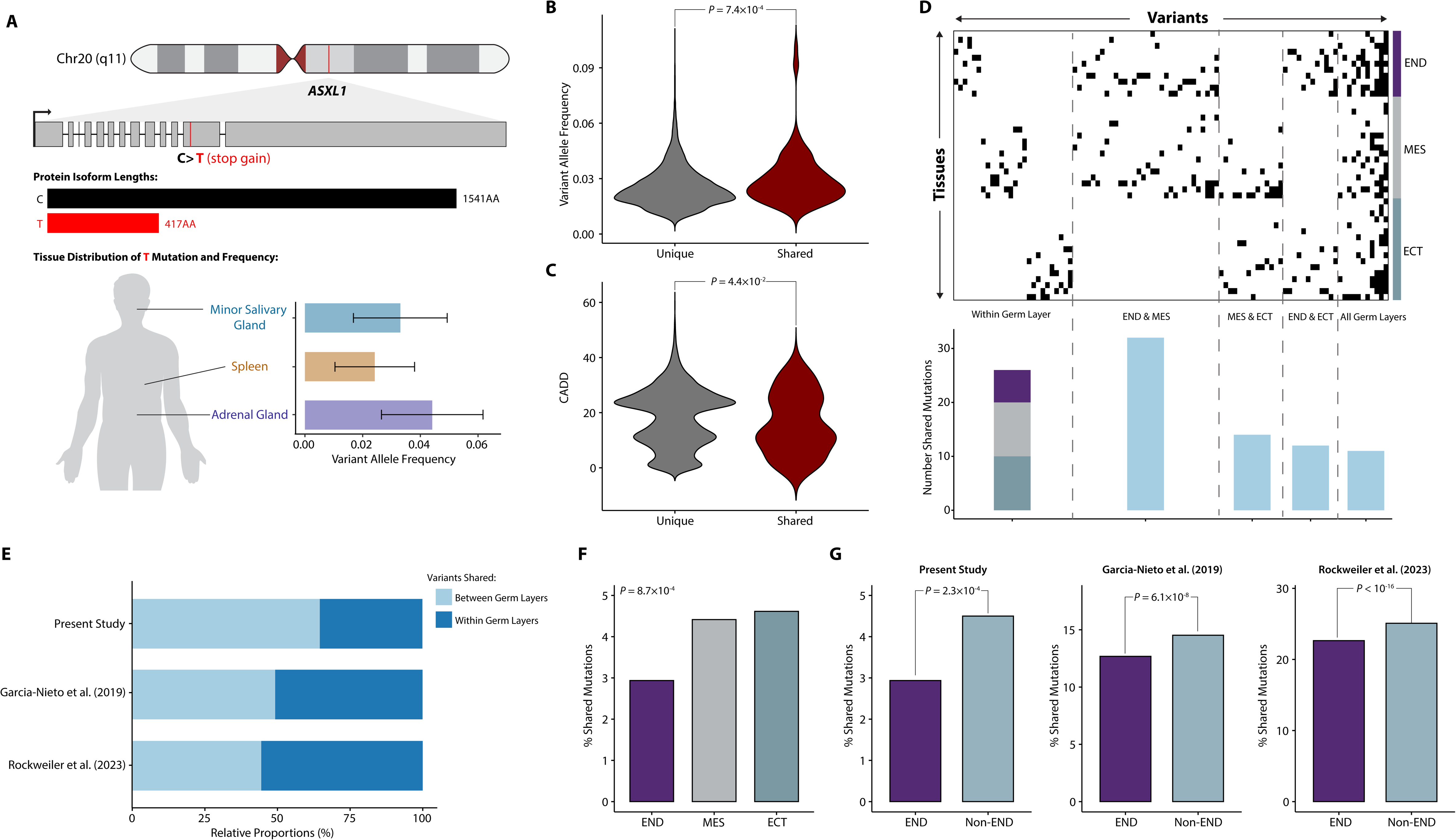
Patterns of shared somatic mutations across tissues. **(A)** Schematic illustration of a shared stop-gain C>T mutation somatic mutation in *ASXL1*. The length of the wild-type (black) and mutated protein (red) is shown. This somatic variant was present in the minor salivary gland, spleen, and adrenal gland from the same donor. **(B)** Distribution of variant allele frequencies (VAF) for shared and unique somatic mutations. **(C)** Distribution of CADD scores for shared and unique somatic mutations. **(D)** Heatmap illustration of all 853 shared somatic mutations (columns) and the tissues they were observed in (rows). Tissues are grouped by germ layer, as indicated on the right side of the heatmap (END = endoderm, MES = mesoderm, and ECT = ectoderm). Variants are sorted based on whether they are shared within one or more germ layer(s) and then by the number of tissues. The barplot below shows the number of shared mutations across different combinations of germ layers. **(E)** The proportion of somatic mutations shared within (dark blue) or between germ layers (light blue). These proportions are shown for our data and two previously published datasets. **(F)** The proportion of shared mutations (relative to all mutations found in tissues from a given germ layer) in each germ germ layers. **(G)** The proportion of shared mutations in the endoderm (END) versus those in non-endoderm (Non-END) germ layers in our study and two previously published data sets. All *P*-values shown were obtained from chi-squared tests.

Consistent with previous findings (32), we reconfirmed a significant positive correlation between VAF and the proportion of donors’ tissues exhibiting shared mutations (Pearson *R* = 0.39; *P* < 10^-16^, Kruskal–Wallis Test; Fig. S2A). No differences of the mutational spectrum between shared and unique SNVs were observed across germ layers (Fig. S2B).

### Most shared somatic mutations are found in tissues that originate from multiple germ layers

The collective patterns of mutation sharing across tissues are depicted in Fig. 4D. On average, shared somatic SNVs were found in 3.4 tissues, with a range of 2 to 27 tissues. Of the shared SNVs, 71.6% (68/95) were identified across two tissues. A mere 4.2% (4/95) were found in ten or more tissues.

We then investigated whether occurrences of shared somatic mutations were more prevalent among tissues originating from the same or distinct germ layers. Notably, only 26 variants were shared among tissues exclusively within a single germ layer (Fig. 4D).

Consequently, the majority (72.6%, 69/95) of shared somatic variants are present in tissues spanning more than one germ layer. A pronounced level of sharing was observed between mesoderm and endoderm compared to other pairwise germ layer comparisons (Fig. 4D).

Furthermore, we identified 11 SNVs shared across tissues derived from three germ layers (Fig. 4D), affecting 3 to 27 tissues from the same donor. For instance, a synonymous variant in the *KEL* gene was detected in the Brain-Caudate (basalganglia) (ectoderm), Adrenal Gland (mesoderm), and Liver (endoderm) of a 66-year-old donor.

We proceeded to conduct a quantitative analysis of shared mutation patterns within and between germ layers. Our focus was on the relationship between the number of tissues exhibiting shared variants and the germ layer patterns from which these tissues originated. As anticipated, SNVs shared across a larger number of tissues were more likely to be found in at least two tissues originating from different germ layers (Fig. S2C). This observation was further reaffirmed by the reanalysis of previously published data from GTEx (30,32), which showed consistent trends (Fig. S2D-E). Considering all shared mutations, 72.6% (69/95) of multi-tissue SNVs are present in at least two tissues from different germ layers. Interestingly, even when somatic variants were found in two tissues, 64.7% (44/68) of shared mutations were discovered in tissues from different germ layers, as opposed to those from the same germ layer (Fig. 4E & Fig. S2C). This proportion, while consistently high, is found to be 49.3% and 44.3% in the studies by García-Nieto et al. (30) and Rockweiler et al. (32), respectively (Fig. 4E; Fig. S2D-E).

### Patterns of shared somatic variants represent developmental bottlenecks

We also sought to determine if the proportion of shared somatic SNVs found in tissues from a specific germ layer varied across different germ layers. Towards this aim, we calculated the proportion of shared mutations relative to all variants found in tissues from a germ layer. We observed a significant difference between germ layers (*P* = 8.7×10^-4^, Chi-squared test; Fig. 4F), primarily driven by a lower proportion of shared variants in the endoderm (*P* = 2.3×10^-4^, Chi-squared Test; Fig. 4G). In line with this, tissues from the endoderm generally exhibited a significantly lower proportion of shared variants compared to those from the mesoderm and ectoderm (*P* = 2.3×10^-2^, Kruskal–Wallis Test; Fig. S2F). Remarkably, upon reanalyzing two additional datasets from GTEx (30,32), we found a similar pattern (*P* = 1.6×10^-19^ & *P* < 10^-16^, Chi-squared Test; Fig. 4H-I). This observation could potentially be attributed to developmental bottlenecks in the endoderm (46,47).

### Rates of somatic mosaicism are highly variable among donors and tissues

To quantify the load of somatic mosaicism, we defined the mutation burden of a sample as the count of somatic variants detected. The normalized somatic mutation burden of a sample was characterized as the mutation burden adjusted by the size of the sample’s exome regions (the number of bases with a minimum of 10× total coverage and a maximum of 5 times the median coverage). The normalized burden of somatic mosaicism exhibits considerable variability across different tissues and donors (Fig. 5A-B). The median normalized mutation load within a tissue varied from 0.21 somatic mutations per Mb in the cerebellar hemisphere to 2.49 somatic SNVs per Mb in the vagina (Fig. 5A). The median normalized burden per donor ranged from 0.42 to 1.45 somatic SNVs per Mb (Fig. 5B). This substantial variability underscores the diverse landscape of somatic variant burdens within and across tissues and donors.

**Figure 5.**
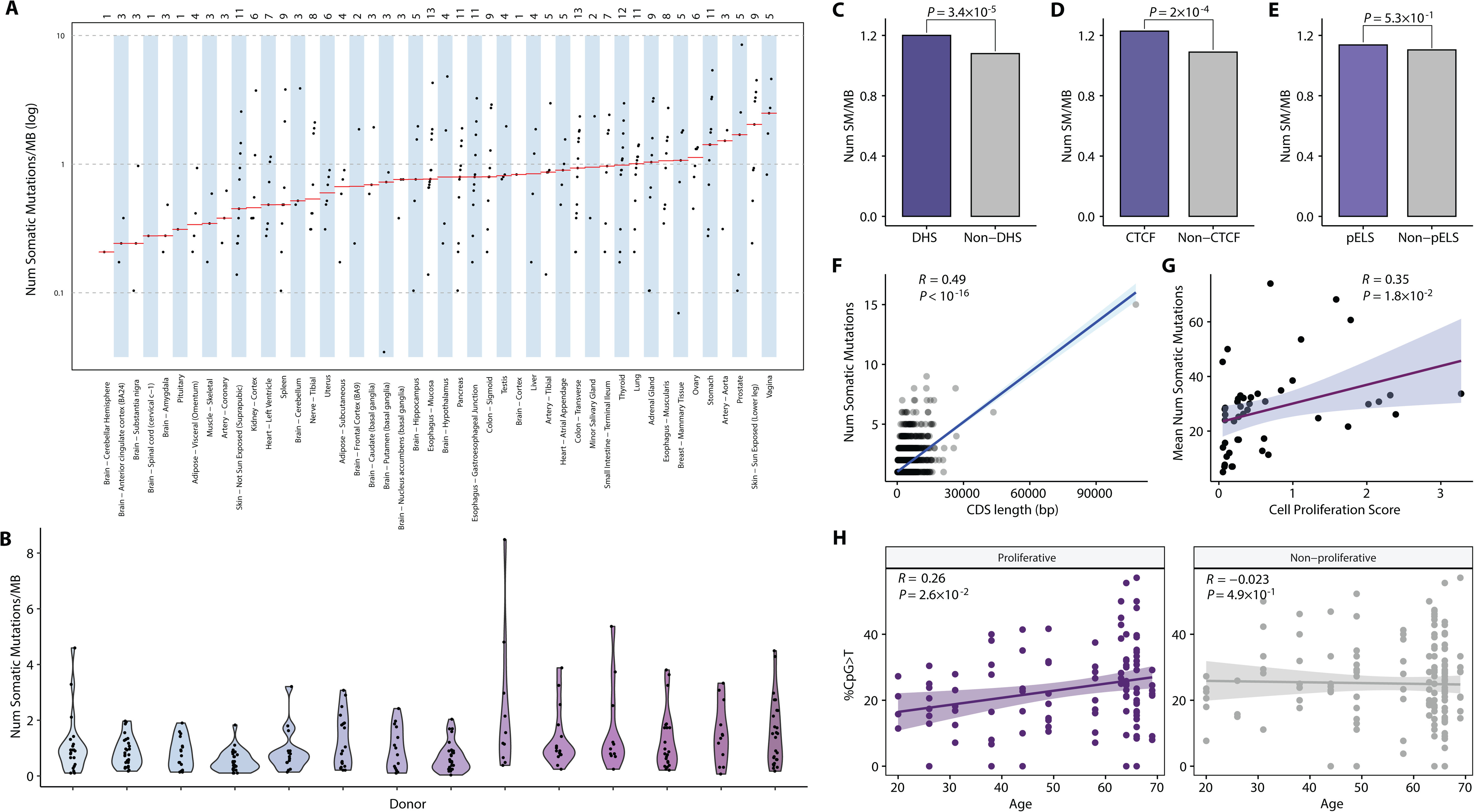
Determinants of somatic mosaicism. **(A)** Variability of the normalized somatic mutation burden (on a logarithmic scale) across tissues. Each dot represents a sample, with the total number of sequenced samples per tissue indicated at the top of the figure. The red solid lines represent median values within a tissue. **(B)** Violin plot showing the distribution of the normalized somatic mutation burden among tissues across the 14 donors. **(C)** Enrichment of somatic mutations in DNaseI Hypersensitive Sites (DHS), (**D**) CTCF sites, and (**E**) proximal enhancer like sequences (pELS). *P*-values were calculated from Chi-squared tests. **(F)** Relationship between CDS length and the number of somatic variants in a gene. **(G)** Correlation between tissue-specific cell proliferation score and the average number of somatic mutations across tissues. In panels (**F**) and (**G**), The Pearson correlation coefficient, *P*-value, and 95% confidence intervals derived from linear regression are shown. **(H)** Relationship between the proportion of CpG>T transitions for each sample and age in proliferative and non-proliferative tissues. The Pearson correlation coefficient, *P*-value derived from a linear mixed model, regression lines, and 95% confidence intervals are shown.

We found that donors and tissues individually account for 15.2% and 17.1% of the variance in the normalized burden, respectively (see STAR Methods). Together, they elucidate 32.3% of the total variance when their effects are considered additively. This combined influence remains relatively stable even after imposing a minimum requirement of five samples per tissue or sequentially eliminating the sample with the highest or lowest normalized burden. Encouragingly, when performing the same ANOVA analysis on a recent GTEx RNA-seq dataset (32), the proportions of variability explained remain strikingly similar, with 17.2%, 20.4% and 37.9% attributed to donors, tissues, and their combined effects, respectively. In alignment with the methodology of Rockweiler et al. (32), we also treated donors as a random effect (STAR Methods). We estimated that donors accounted for 9.2% of the variability, a figure nearly identical to that reported in their study (32), 8.8%.

### Factors associated with the burden of somatic mutations

We investigated the relationship between a number of factors and rates of somatic mutations. It is well known that chromatin structure is associated with rates of germline mutations and somatic mutations in cancer (48). However, contrary to cancer studies, which typically observe a relative decrease in mutation density within DNase I hypersensitive sites (DHS) (48), we found a significant enrichment of somatic variants within these DHSs (*P* = 3.4×10^-5^, Chi-squared test; Fig. 5C). This observation is consistent with simulations predicting a higher relative density of mutations in DHSs based on sequence context (48).

This phenomenon could potentially be explained by the increased accessibility of DNA at DHSs, which may render it more prone to damage compared to less accessible DNA located elsewhere. Interestingly, these enrichments were more specifically linked to CTCF regions (*P* = 2.0×10^-4^, Chi-squared test; Fig. 5D), rather than proximal enhancer like sequencers (pELS, *P* = 0.53, Chi-squared test; Fig. 5E). CTCF is an insulator-binding protein and is involved in chromosome 3D structure. Mutations at CTCF sites might prevent normal CTCF binding and result in different topologically associated domains (68). pELS are candidate enhancers within 2kb of transcription start sites.

Next, we tested the relationship between CDS length and the number of somatic mutations observed in genes. The observed Pearson correlation coefficient was 0.49 (Fig. 5F), implying that 24% of the variation in the gene-based somatic burden can be explained by CDS length. Interestingly, for germline mutations from the gnomAD database (49), notably higher Pearson correlation coefficients are observed between CDS length and number of variants. Specifically, the Pearson correlation coefficient for rare (≤1%) and common (>1%) GnomAD variants are 0.88 and 0.6 (Fig. S3A-B), respectively.

### The burden of somatic mosaicism exhibits a positive correlation with tissue-specific rates of cell proliferation

We confirmed a positive linear correlation between tissue-specific cell proliferation (as determined by MKI67 expression, a marker of proliferation) (31) and the average number of accumulated distinct mutations (Pearson *R* = 0.35; *P* = 1.8×10^-2^, Linear Regression; Fig. 5G) or average normalized overall burden (Pearson *R* = 0.34; *P* = 2.1×10^-2^, Linear Regression; Fig. S3C) across various tissues. This finding is consistent with the outcomes reported by Yizhak et al. (2019) (31), and is further corroborated by the reevaluation of data from two additional GTEx studies (30,32) (Fig. S3D-E). Furthermore, this significant correlation is attributed to C>T transitions (Pearson *R* = 0.39; *P* = 7.2×10^-3^, Linear Regression; Fig. S3F). Reanalysis of two previous data sets also reveal significant positive correlations between the burden of C>T transitions and cell proliferation (Fig. S3G-H).

### The interplay of CpG>T mutations, age and cell proliferation

To investigate effects of aging on somatic mutations, we employed a linear mixed model that related both biological and technical variables to the proportions of CpG>T mutations at the sample level (STAR Methods). The proportions of CpG>T transitions for each sample acted as an estimate of the observed aging signature (SBS1) (9), normalized by the count of detected somatic mutations, thereby accounting for the variability in sequencing coverage across samples. This model allowed for random slopes to vary among donors, as we anticipated the mutation rate to be donor-dependent. Age showed marginal significance (*P* = 3×10^-2^, Linear Mixed Model), while the effect of tissues was more pronounced (*P* = 1.6×10^-2^, Linear Mixed Model). However, the significance of age (*P* = 0.17) vanished when tissues were not accounted for in the model. This led us to hypothesize interplays between CpG>T transitions, age, and tissue types, particularly whether tissues were proliferative or not.

To disentangle this combined influence, we integrated tissue-specific cell proliferation and examined its interactions with aging and CpG>T mutations (STAR Methods). We classified tissues with the top 30% cell proliferation scores as proliferative, and the remaining tissues as non-proliferative. We extended previous analysis (31) and revealed significant positive associations between age and the proportions of CpG>T mutations (Pearson *R* = 0.26; *P* = 2.5×10^-2^, Linear Mixed Model; Fig. 5H) exclusively within proliferative tissues. In contrast, samples from non-proliferative tissues showed no significant relationship with age (*P* = 0.49, Linear Mixed Model; Fig. 5H). Importantly, these findings persisted even after (i) allowing for random slopes to vary between donors (shifting from a Linear Mixed Model to standard linear regression), (ii) adjusting the cell proliferation score cutoff to different deciles (Fig. S3I), and (iii) dividing all individuals into two distinct age groups (Fig. S3J). In contrast, no such correlation was observed between aging and the proportions of non-CpG C>T transitions, even when considering cell proliferation (Fig. S3K).

Note that upon substituting tissues with cell proliferation scores in either the linear mixed model or standard linear regression, we failed to detect significant signals associated with age or cell proliferation. Consequently, we inferred two possibilities: Firstly, the statistical power is diminished when cell proliferation is incorporated into the model, possibly due to the noisy measurements of expression markers in tissues or the nonlinear relationships between these markers and the proportions of CpG C>T transitions. Secondly, we speculated the existence of additional factors, beyond cell proliferation, that contribute to the intricate interplay among CpG>T transitions, age, and tissue types.

### Characteristics of clinically relevant somatic mutations

To quantify how often pathogenic variants occur in apparently healthy individuals, we defined Clinically Relevant Mutations (CRM) as those somatic mutations that are present in the Human Gene Mutation Database (HGMD) (50), ClinVar (51), and two Cancer databases (Database of Curated Mutations and Cancer Genome Interpreter) (52,53) (see STAR Methods). In total, 0.7% (54/8243) of all somatic SNVs are classified as clinically relevant, with one variant appearing in two tissues of the same donor and the remaining 53 being distinct. Specifically, 45, 26, and 4 somatic variants are documented in the HGMD, ClinVar, and cancer databases, respectively (Fig. 6A).

**Figure 6.**
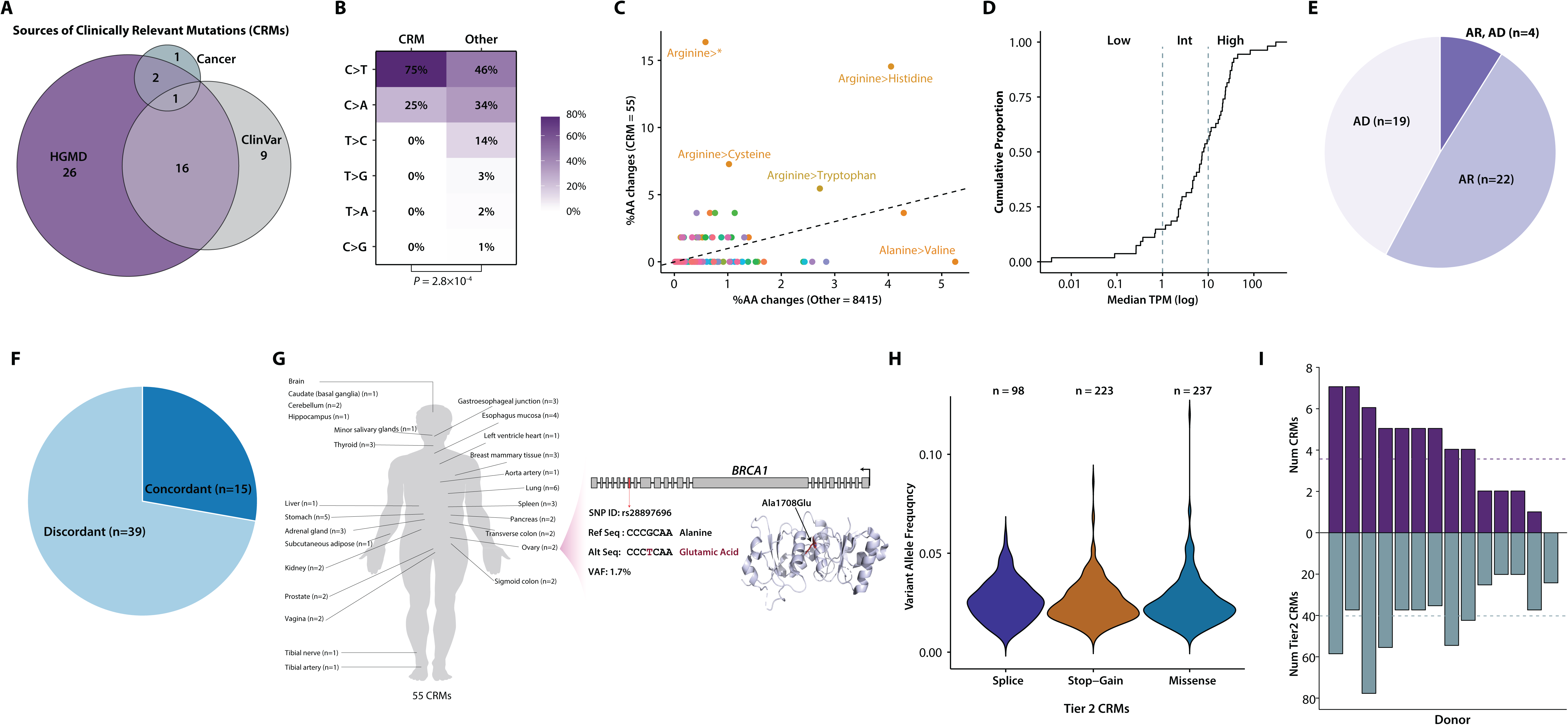
The burden of clinically relevant somatic variants among individuals. **(A)** The number of putative clinically relevant somatic mutations (CRMs) identified in HGMD, ClinVar, and cancer databases (DoCM and CGI). **(B)** Comparison of the mutational spectrum between CRMs and other mutations. *P*-value is calculated from a Fisher’s Exact Test. **(C)** Contrast of the proportions of amino acid alterations in CRMs versus other mutations. A dashed line represents equality (y=x). A spectrum of amino acid changes are color-coded. Stop-gained variants are denoted as *. **(D)** Empirical cumulative distribution function (eCDF) of median gene-level TPM by tissue in RNA-seq (on a logarithmic scale). Genes are classified into low-, intermediate-, and high-expression categories based on cutoffs of 1 and 10. **(E)** Pie chart portraying the distribution of inheritance modes: AD, AR, and a blend of AR and AD. **(F)** Pie chart indicating the concordance between the tissues where CRMs were discovered and the tissues impacted by the phenotype. **(G)** Schematic representation of 25 tissues identified with 55 observed CRMs. Numbers in brackets indicate the count of CRMs detected in each tissue. An example of a CRM in *BRCA1* detected in the ovary is provided, showcasing gene and protein structure alongside fundamental CRM information (the nucleotide alteration, amino acid change, 3bp-flanking sequence, VAF and median TPM of *BRCA1* in the ovary). **(H)** Tally of detected somatic mutations across a range of clinically relevant gene sets. **(I)** Enumeration of CRMs (top) and somatic mutations falling within an expansive definition of clinically relevant gene sets (bottom) across individuals. Dashed lines represent the average values.

CRMs exhibit a unique mutational spectrum compared to non-CRMs (*P* = 2.8×10^-4^, Fisher Exact Test; Fig. 6B). They show a preponderance of C>T transitions and a decrease in all other types (Fig. 6B), consistent with previous findings (54). CRMs are predominantly characterized by mutated Arginine, particularly leading to stop gain (16.4%), Histidine (14.5%), Cysteine (7.3%), and Tryptophan (5.5%) (Fig. 6C). Conversely, CRMs are depleted of Alanine-to-Valine changes (0%) compared to other variants (5.3%) (Fig. 6C).

We examined the association of CRMs with gene expression levels in RNA-seq data. Genes were stratified into low-, intermediate-, or high-expressed categories using cutoffs of 1 and 10. Among the 54 CRMs, the majority are in genes classified as intermediate (40.7%) or high (44.4%) expression, with only 14.8% in low-expressed genes (Fig. 6D). Furthermore, among the 45 CRMs with identified inheritance modes, 42.2% (19/45) are autosomal dominant (AD), while 48.9% (22/45) are autosomal recessive (AR) (Fig. 6E). An additional 8.9% (4/45) of CRMs exhibit patterns of both AR and AD inheritance (Fig. 6E).

We further explored whether tissues harboring CRMs align with those affected by associated phenotypes. 27.8% (15/54) of tissues with CRMs show consistency with tissues affected by phenotypes in the databases (Fig. 6F). Another 72.2% (39/54) of CRMs occurred in tissues that were not directly affected by associated phenotypes (Fig. 6F). However, altered expression levels or other deleterious molecular phenotypes in unrelated tissues may be induced by disease-causing somatic alleles (4,55,56).

Twenty five out of 46 tissues were found with at least one CRM (Fig. 6G). The three tissues with the highest number of CRMs are Lung, Stomach, and Esophagus-Mucosa, with 6, 5, and 4 detected CRMs, respectively. An example of a missense CRM in the 18th exon of *BRCA1* is provided (Fig. 6G). This CRM in *BRCA1*, with a VAF of 1.7%, was observed in the ovary of a 64-year-old donor. The 1708th residual was mutated to Glutamic Acid (Glu) from Alanine (Ala). *BRCA1* is intermediately expressed in the ovary, given that its median transcript per million (TPM) in the GTEx tissues is 1.54 (29).

### Expanding the concept of CRMs and estimates of donor-specific burdens

When applying a broader concept of clinically relevant genes (STAR Methods), we observed somatic SNVs falling into varied sets of functionally important genes (Fig. 6H), ranging from 3,616 HGMD disease-causing mutations (50) to 97 cancer mutations curated by DoCM (52).

Finally, we investigated the donor-specific burdens of CRMs and somatic mutations falling into diverse functionally important gene sets. 13 out of 14 donors are identified with at least one CRM (Fig. 6I top). On average, each donor harbored 3.93 CRMs (Fig. 6I top). At maximum, two donors carried seven CRMs (Fig. 6I top). There are on average 323.4 somatic mutations in functionally important genes for each donor, ranging from 197 to 605 somatic variants (Fig. 6I bottom). This underscores the importance of studying somatic mutations in personal genomics.

## Discussion

In our study, we conducted high-coverage exome sequencing on 265 GTEx samples, encompassing 46 unique tissues from 14 individuals. Utilizing mSOMA, a novel method that is tailored to the structure of our data, we were able to detect somatic mutations directly from bulk DNA. Our results offer new insights into shared multi-tissue somatic mutations, functionally significant mutations, and clinically relevant variants. These findings illuminate the landscape and characteristics of somatic mosaicism in healthy individuals, setting the stage for future research investigations and potential clinical implications.

Bulk DNA sequencing facilitates the discovery and identification of somatic mosaicism both within and among healthy individuals. It presents a clinically viable and effective approach with the potential for scalability. However, one limitation of our study is that bulk DNA sequencing is more adept at detecting common somatic variants, as opposed to the rarer forms of mosaicism. These rarer forms could potentially be identified using techniques such as laser capture microdissection, duplex sequencing, or single-cell sequencing.

We observed heterogeneity between our cohort and other GTEx studies. For instance, we identified some somatic mutations in our samples that were absent in the RNA-seq data and vice versa. Donors and tissues combined account for 32.3% of the variance in the normalized burden in our data, while such an estimated proportion is 37.9% in Rockweiler et al. (2023) (32). These differences could potentially be attributed to the actual cells sampled for each tissue in DNA versus RNA, varying methods in calling somatic variants, and differences in sample sizes. Despite the more straightforward approach of studying somatic mosaicism in DNA versus RNA, we observed a similar overall trend and drew parallel conclusions between DNA and RNA data. For example, we noted a concordant positive correlation between somatic variant load and tissue-specific cell proliferation, and a consistently significantly lower proportion of shared variants in tissues deriving from the endoderm compared to those from the mesoderm and ectoderm. Both our study and Rockweiler et al. (2023) (32) predicted approximately 2-3% higher variability explained by tissues than donors.

Through our simulations, we found that the distributions of the expected and observed ratios of missense to synonymous somatic mutations differ from those of germline variants. This discrepancy could be attributed to the varying DNA repair pathways active in different tissues and the sensitivity of these tissues to environmental pressures. Furthermore, we noted that the empirical ratios of missense to synonymous somatic mutations exhibit variation across tissues (Fig. S4A) and across different studies (Fig. S4B).

In our cohort, we found no direct evidence of positive selection on cancer genes in normal tissues. This is primarily because we did not observe elevated VAFs in unique variants present in oncogenes or tumor suppressor genes (TSG) (Fig. S4C). This lack of evidence could be attributed to the small sample size of our study, particularly given that 43% (6/14) of the participants were under the age of 50. An alternative method for investigating positive selection in normal tissues involves the refinement of dN/dS, taking into account the variability of the mutation rate across the human genome (44). However, it’s important to note that complications have been identified in using dN/dS for genetic sequences sampled from a single population (57).

## STAR * METHODS

### KEY RESOURCES TABLE

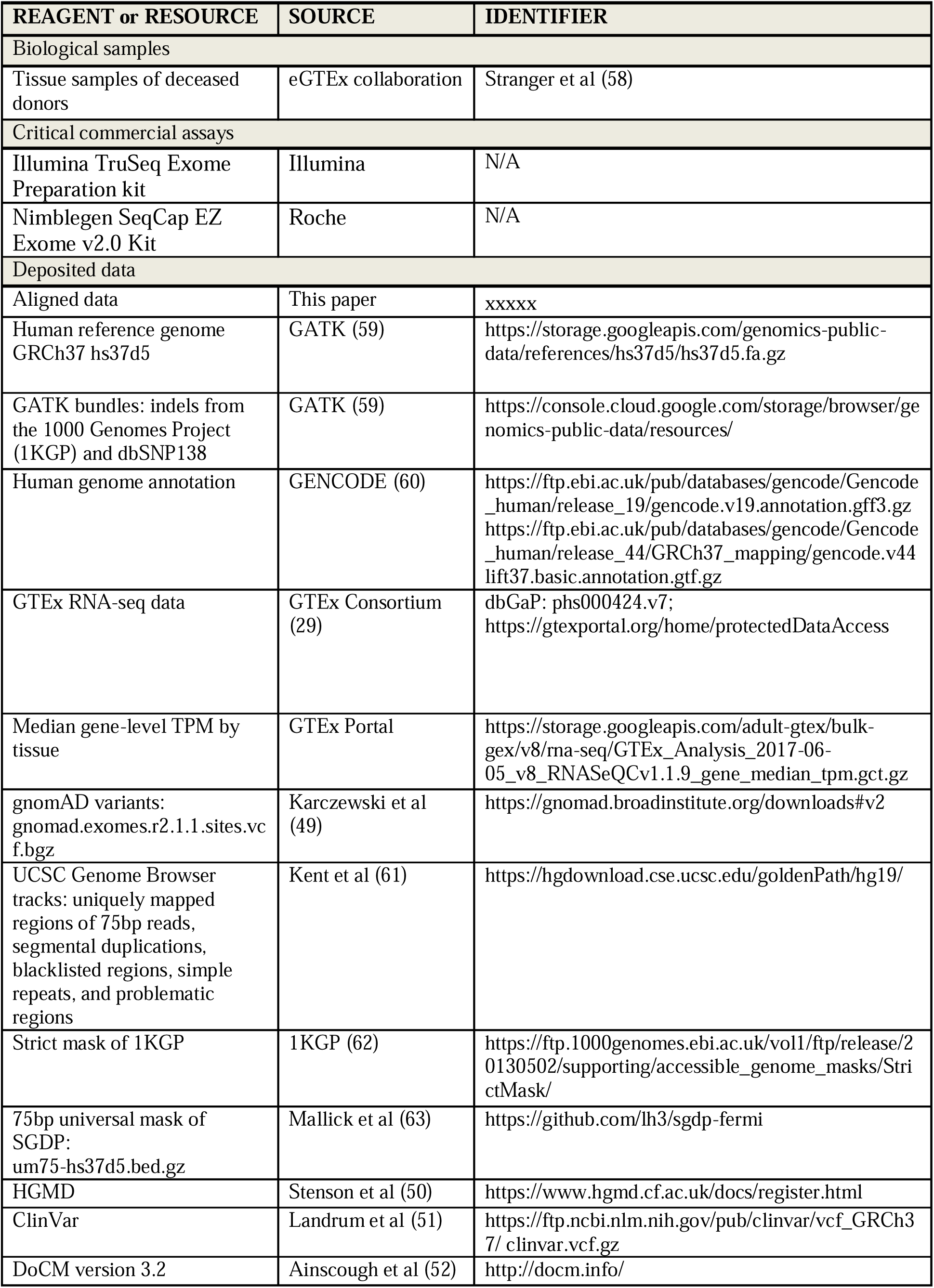
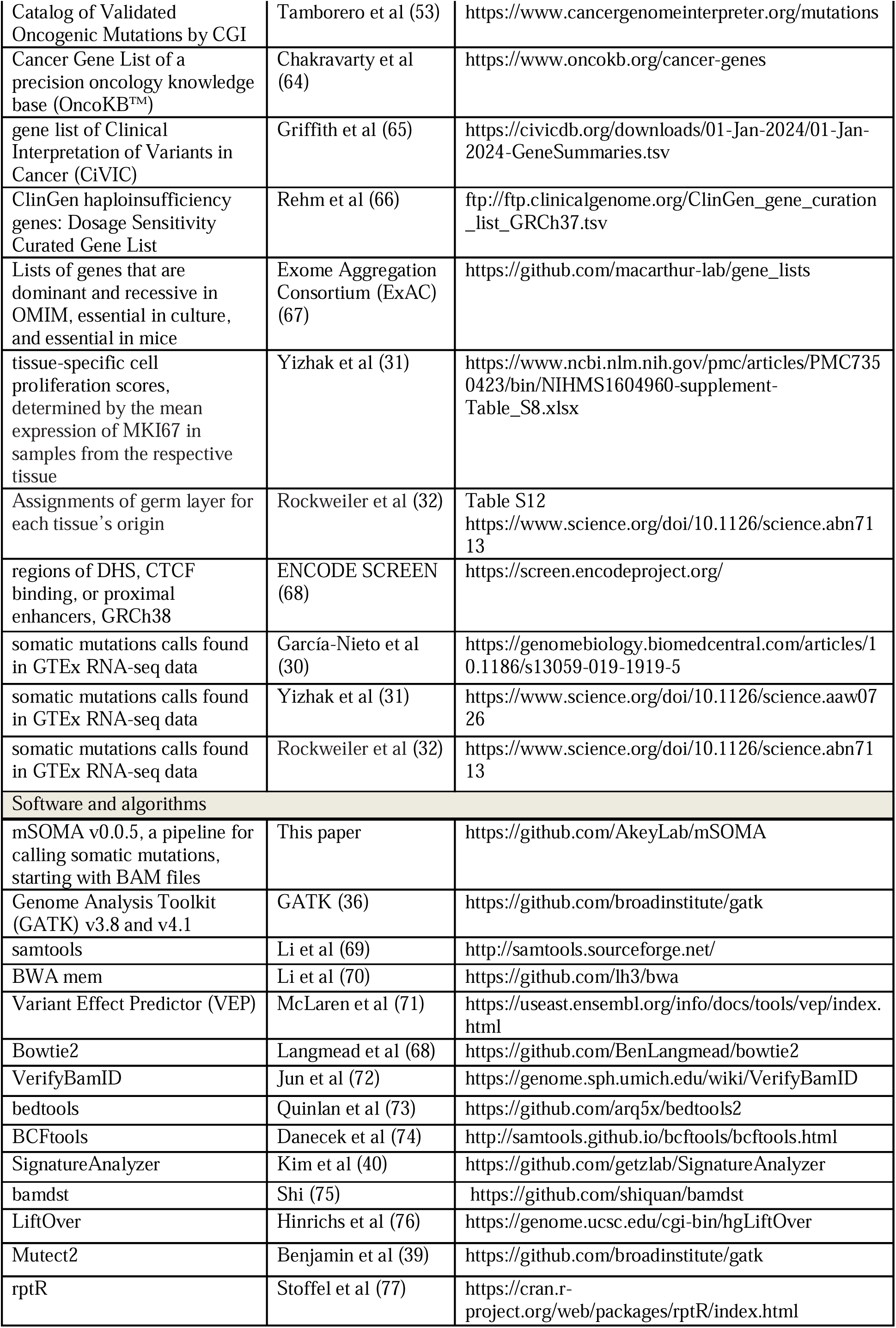

### RESOURCE AVAILABILITY

#### Lead contact

Further information and requests for resources should be directed to and will be fulfilled by the lead contact, Joshua M. Akey.

#### Materials availability

This study did not generate new unique reagents.

#### Data and code availability

All protected mapped data of 265 GTEx samples have been deposited at the database of Genotypes and Phenotypes (dbGaP) and are publicly available as of the date of publication. The accession number is listed in the key resources table. mSOMA is open-source and publicly available at https://github.com/AkeyLab/mSOMA and is provided as an executable Docker image to facilitate ease of use and reproducibility. Any additional information required to reanalyze the data reported in this work paper is available from the lead contact upon request.

### EXPERIMENTAL MODEL AND STUDY PARTICIPANT DETAILS

#### GTEx donors and tissue samples

Detailed information about the process of donor enrollment, consent, and biospecimen collection can be found in previous publications (29,78,79). In brief, the human donors involved in this study were deceased individuals. Tissues were sourced from either post-mortem donors or organ donors. Consent for participation in the study, which included the collection and de-identification of tissue samples, was obtained from the immediate family members of the donors.

We collected DNA from 14 donors of the Genotype-Tissue Expression (GTEx) project, covering 46 different tissues. This was part of the enhanced GTEx Consortium (eGTEx) initiative, which aimed to augment existing gene expression data with additional molecular phenotypes for a subset of GTEx donors (33). The 46 tissues encompassed a broad spectrum of organ and developmental lineages. The sampled tissues include adipose tissue, adrenal gland, coronary artery, brain, transverse and sigmoid colon, esophagus, heart, kidney, liver, lung, minor salivary gland, skeletal muscle, nerve, ovary, pancreas, pituitary, prostate, epithelial skin tissue exposed and unexposed to the sun, ileum (Peyer’s patch), spleen, stomach, testis, thyroid, uterus, and vagina. Additional details regarding the 14 analyzed donors are presented in Table S1.

#### DNA extraction, exome capture, library preparation, exome capture and sequencing

A total of 299 unique samples, along with 4 replicates, were subjected to high-coverage exome sequencing at the Northwest Genomics Center (NWGC) at the University of Washington. For the majority of DNA samples, the input ranged between 150 and 200ng. Libraries were prepared using the Illumina TruSeq Exome Preparation kit. The amplified libraries were then captured using the Roche Nimblegen V2 Exome, in accordance with the manufacturer’s instructions. Sequencing was performed on the Illumina HiSeq 4000, generating paired-end reads of 75bp.

### METHOD DETAILS

#### Read alignment and standard GATK Best Practice

Adapters were marked using Genome Analysis Toolkit (GATK, 4.1.7.0 unless otherwise noted) (59) MarkIlluminaAdapters. Following this, sequencing reads were aligned to the human genome reference GRCh37 hs37d5 (59) using the mem function of the Burrows-Wheeler Aligner (BWA, 0.7.17-r1188) (70) on a per lane basis. The lane-level BAMs were subsequently merged into a single BAM for each sample using GATK MergeSamFiles. PCR duplicate reads were identified and labeled within each BAM using GATK MarkDuplicates. A list of target intervals for indels was generated for each donor using the GATK RealignerTargetCreator (3.8-1-0-gf15c1c3ef). This list included potential indels observed in samples from the specific donor and standard resources of GATK bundles (59), which are known indels from 1KGP and dbSNP138. Local realignment around these candidate Indels was carried out independently for each sample using GATK IndelRealigner (3.8-1-0-gf15c1c3ef). Base quality scores were recalibrated using GATK BaseRecalibrator and ApplyBQSR, with the updated base quality scores retained in the BAMs. The final BAM files were stored for subsequent quality control, variant calling, and other downstream analyses.

#### BAM-based sample-level quality control

We implemented a rigorous quality control process at the sample level using BAM files. This process was designed to detect sample swaps, mislabels, cross-sample contamination, and discrepancies between reported and genetically determined sex, leading to a refined cohort. Firstly, we used GATK CrosscheckFingerprints and ClusterCrosscheckMetrics to identify instances where sample BAMs were incorrectly linked to a donor, which might be due to sample swaps or mislabels. Samples flagged as UNEXPECTED_MISMATCH were considered mismatched with the donor, resulting in the exclusion of 15 samples. Secondly, we identified discrepancies between the reported and genetically determined sex of a sample. This was based on the coverage of Chromosome Y, normalized to the coverage and size of capture regions on Chromosome X, calculated as:

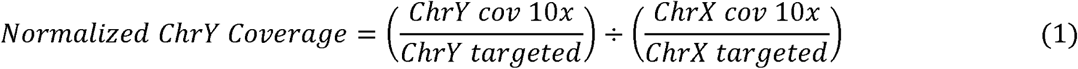

Only non-PAR regions of sex chromosomes were used in sex classification. Samples with normalized Y coverage greater than 0.8 were classified as male, and those with normalized Y coverage less than 0.2 were classified as female. Six samples, including four that were already identified as mislabeled, were excluded due to ambiguity or inconsistency between reported and genetic sex.

Lastly, we applied VerifyBAMID (v1.1.3) (72) software to detect potential cross-sample contamination. This was based on a set of approximately 200,000 Axiom Exome Plus sites obtained from the GATK bundle resource. We conservatively identified and excluded 16 samples that were potentially contaminated, with a FREEMIX score greater than 0.01. Following the sample quality control process, we were left with 265 unique samples and 3 replicates. We generated BAM-based mapping metrics using bamdst (v1.0.9) (75). The average sequencing coverage across target regions is presented in Fig S1A.

#### Pre-filters of reads and bases prior to somatic SNV calling

After the mapping process, we applied a series of pre-filters to the reads and bases before calling somatic SNVs. We started by trimming 5 base pairs from both ends of the reads. Then, we utilized samtools (69) to filter the reads according to various criteria. Specifically, we discarded reads if their mapping quality was below 20 or if they were flagged as “UNMAP” (segment unmapped), “SECONDARY” (secondary alignment), “QCFAIL” (not passing quality controls), “DUP” (PCR or optical duplicate), “SUPPLEMENTARY” (supplementary alignment), or “IMPROPER PAIR” (each segment improperly aligned according to the aligner). We also filtered out reads if any Indels were detected, if more than 10% of bases were mismatched, or if more than 50% of the bases were soft-clipped. Lastly, we retained only those bases with a base quality score of at least 20 for further analysis.

#### Definition of callable regions

To mitigate alignment errors and other technical artifacts from short pair-end reads, we used bedtools (73) to establish a callable region for somatic SNV identification. We focused on autosomes and excluded sites not on target regions or within a 10 base pair padding. We systematically filtered out sites falling into various categories, including regions within 5bps proximal to Indels identified in the 1KGP (62), low complexity regions, HLA loci in Chromosome 6, centromeric regions, segmental duplications (61), simple repeats (61), blacklisted regions (61), and problematic regions (61). We also applied the strict accessibility mask from 1KGP (62) and the universal mask from SGDP (63).

We included only germline reference homozygotes for each donor. To obtain these, we used the GATK HaplotypeCaller (with ploidy set to 2), aggregated sample-level GVCFs per donor, and performed GATK (version 4.2.0.0) GenomicsDBImport and GenotypeGVCFs at all sites within the targeted regions of all autosomes. If the depth of coverage (DP) was less than or equal to 10, we set the genotype of a tissue sample to missing. We defined germline reference homozygotes for each donor as sites where at least 95% of tissues from the same donor were called as reference homozygotes.

Finally, we derived sample-specific callable regions by: 1) including sites where a tissue sample was called as a reference homozygote with at least 10 reads covered, 2) excluding loci with sequencing coverage below 10x or exceeding 5 times the median coverage for each sample, and 3) including loci with at most two observed base calls for each sample. This systematic approach ensured the reliability and accuracy of our subsequent analyses.

#### Developing a Beta-binomial model for single-sample somatic SNV calling

In samples of bulk sequencing, somatic variants often present in low variant allele frequencies (VAFs), and sequencing errors can further confound the analysis. To distinguish somatic mutations from various artifacts, we expanded the application of the Beta-binomial model. This model, which has been previously used to characterize distributions of sequencing errors (80,81) or RNA-seq counts (82), was applied to identify potential somatic mutations. Briefly, we modeled alternative allele counts locally by Beta-binomial distributions. True positive somatic mutations, which are distinguished by a higher count of variant-supporting reads compared to the local error rate, were identified through hypothesis testing.

It is important to note that we focused on the trinucleotide context, a small-scale feature of the genomic architecture that impacts the substitution rate (83). The trinucleotide context (“3-mer”) takes into account single 5′ and 3′ nucleotides that flank the polymorphic middle position. There are six classes of base substitution, each referred to by the pyrimidine of the mutated Watson-Crick pair: C>A, C>G, C>T, T>A, T>C, T>G (all referred to by the pyrimidine of the mutated Watson-Crick pair). By combining four possible reference bases for each independent flanking position, this trinucleotide sequence context model results in a total of 96 (4 × 6 × 4) possible substitution types.

Using samtools (69) mpileup, we obtained allele counts across callable regions. Based upon this information, we performed single-sample variant calling based on trinucleotide context using the Beta-binomial model. Specifically, independently for trinucleotide context for each sample, counts of alternative (non-reference) alleles (*AC_i_*) at the (germline) homozygous-reference site *i* are assumed to follow a Binomial distribution with parameter *p^j^* (error rate) under a trinucleotide sequence context *j* (*j*=1,2,…,96), which is a random variable that follows a Beta distribution with parameters α*^j^* and β*^j^* (Formula 2-4).

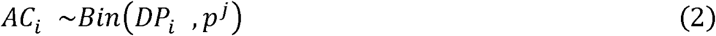

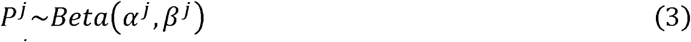

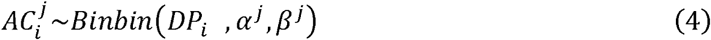

Somatic SNV calling was framed as a hypothesis testing problem. The null hypothesis posited that observed additional non-reference alleles stemmed from sequencing or other technical errors, while the alternative hypothesis suggested somatic mutations. The test statistic, the observed alternative allele count (*AC_i_*), followed a Beta-binomial distribution under the null hypothesis, calculated independently for contexts and samples. Calls with a *P*- value less than 10^-8^ were deemed candidate somatic variants.

#### Calling candidate somatic mutations with mSOMA

We utilized mSOMA version 0.0.5 to generate Betabinomial *P*-values, initiating the process with BAM files. The procedure encompassed two steps: 1) pre-calling filters and conversion of BAM files into tabulated locus-based allele count information, and 2) execution of initial variant calling by procuring maximum likelihood estimates (MLE) of the context-based Betabinomial parameters and calculating locus-level *P*-values accordingly. These two steps were accomplished using the following commands and parameters:

~~~
betabin count \
 --bed $sample.callable.region.bed \
 --fasta hs37d5.fasta \
 --min-MQ 20 \
 --require-flags 2 \
 --exclude-flags 3844 \
 --max-indel 0 \
 --mismatch-frac 0.1 \
 --softclip-frac 0.5 \
 --seq-length 75 \
 --ntrim 5 \
 --min-BQ 20 \
 --min-depth 10 \
 --max-alt-allele 1 \
 --output $sample.counts.gz \
 $sample.bam
betabin mle \
 --output $sample.pvals \
 --ab $sample.ab \
 --min-depth 10 \
$sample.counts
~~~

#### Post-filters applied to remove artifacts

To enhance the robustness and precision of our variant calling, we employed an extensive post-filtering process. This process is designed to mitigate various artifacts that could potentially hinder the accurate identification of somatic SNVs. We discarded candidate variants with a *P*-value less than 0.01 from a Fisher’s Exact test, which compares the strand bias of bases supporting the reference and alternate allele. We also eliminated candidate mutations with an alternative allele count of less than 3 and those with a VAF of less than

0.005. Variants within 5 base pairs of germline variants of the same donor were removed, with germline variants identified using GATK HaplotypeCaller with a ploidy set to 2. We filtered out loci found with gnomAD (49) variants (SNVs or Indels) with an AF greater than 0.01%. To further refine our results, we applied a reference leakage filter as suggested by Yizhak et al (31), which removed candidate variants identified with a sequence of five or more identical reference bases either upstream or downstream of the variant itself. To exclude potential hotspot mutations, we discarded variants with a minimum distance to other calls of the same sample of less than 20 base pairs. Calls made in more than one individual with a Beta-binomial *P*-value less than 10^-8^ were filtered out. To minimize mapping biases introduced by BWA (70), we included only calls with at least one read covering the same variant allele using Bowtie2 (84) as an additional mapper. Lastly, to control for potential contamination across donors, we removed genomic locations that overlap with germline variants found in any donor.

#### Evaluating power and FDR of mSOMA through simulation

To assess the power and FDR of mSOMA in detecting somatic SNVs, we conducted a simulation study. This study involved 1480 mixed samples, derived from reference and alternative allele counts of our eGTEx sequencing data. The somatic mutation frequency ranged from 1% to 10% in increments of 0.25%. We specified coverage depths to be 50, 100, 300, and 500x. For each combination of somatic mutation frequency and sequencing coverage, we performed ten iterations.

Initially, we used the GATK HaplotypeCaller to determine germline reference and alternative homozygotes for each donor. A locus was classified as such if 95% of tissues from the same donor were labeled as “0/0” or “1/1”, respectively. To create a mixed sample, we selected two random tissue samples from two unrelated individuals, one as a major sample and the other as a minor sample. The minor sample, with germline alternative homozygotes, served as the source of mimic somatic mutations. This process was repeated for ten pairs of random major and minor samples.

To evaluate the impact of coverage on somatic variant calling, simulated data were generated with per-locus depths (i.e., the sum of reference and alternative allele counts) of 50, 100, 300, and 500X. To emulate a mixed cell population, we obtained per-locus counts of reference and alternative alleles as a weighted average from the depth information of two samples and expected somatic mutation frequency. The simulated allele counts were analyzed using the Beta-binomial model of mSOMA, without specifying contexts.

In the simulated sample, true somatic mutations were identified as sites exhibiting reference homozygotes in the major individual and alternative homozygotes in the minor individual (i.e., AA in Individual 1 and GG in Individual 2). Conversely, sites displaying reference homozygotes in both individuals were deemed non-somatic sites. This simulation strategy allowed for the assessment of detection accuracy, including power and false discovery rate (FDR), as a function of coverage and somatic mutation frequency.

#### Assessing the reproducibility of somatic mutation calls with three pairs of duplicate samples

As part of validation, we sequenced three sets of replicate tissue samples utilizing identical DNA libraries to scrutinize the reproducibility of somatic mutation calls. Please note that duplicate samples of the esophagus and colon were obtained from one individual, while duplicate samples of the small intestine were procured from a different, unrelated individual. For each pair, one duplicate was randomly chosen for somatic mutation discovery using mSOMA, while the other served as a validation sample to confirm the identified somatic mutation calls.

We evaluated the concordance between the discovery and validation duplicates by computing the replication rate. This rate is defined as the percentage of mutation calls in the discovery sample that were supported by at least one high-quality read with the same variant allele in the validation replicate. To enhance the precision of our results, we replaced three discovery samples in a cohort of 265 samples with their corresponding validation samples and applied additional post-filters to obtain refined calls for subsequent analyses in Fig. S1B.

#### Performance comparisons of mSOMA and Mutect2 using colon samples

To evaluate the efficacy of different somatic SNV callers, we applied mSOMA to analyze ten samples of Transverse Colons and juxtaposed the results with those obtained from Mutect2 (36). In the absence of a distinct test-reference structure within our dataset, and lacking an apparent reference sample, we explored the influence of reference sample selection on somatic variant identification using Mutect2. In this context, we designated esophagus and stomach tissues from the same donors as matched-normal references.

Subsequently, we independently applied post-filters specific to mSOMA and Mutect2 to derive the final sets of calls for comparison.

#### Validation rates of somatic mutations identified in DNA-seq samples via RNA-seq

To enhance the accuracy of our somatic mutation detection from bulk DNA sequencing, we undertook a comparative analysis with RNA sequencing data. We procured mapped BAM files of RNA-sequencing reads from the dbGAP GTEx project (29) version 7, which were aligned to the human genome reference GRCh37. In an effort to validate our overall SNV calling pipeline, we paired bulk DNA-seq and exome RNA-seq samples from identical tissues and individuals. Of the cohort studied, 80.7% (214/265) of the DNA-seq samples had corresponding RNA-seq samples. We utilized samtools (69) mpileup to obtain allele counts of the mutations discovered in the RNA-seq samples.

Subsequently, we investigated the proportion of mutation calls from bulk DNA-seq that had supporting reads for the same alternate allele in the matched RNA-seq sample. A somatic variant detected in DNA-seq was considered validated if at least one identical variant allele was present in the corresponding RNA-seq sample. The validation rate was thus determined as the percentage of DNA somatic mutations that were corroborated using RNA-seq. This approach provides a robust measure of the reliability of our somatic mutation calls.

#### Annotation of variants and genes

After quality control, somatic variants were annotated using Variant Effect Predictor (VEP) (71). Functional classes of variants are manually reviewed and determined if they are annotated with more than one functional class. Somatic mutations were classified as deleterious if their CADD scores (43) were 30 or higher.

Somatic variants are deemed as clinically relevant if they meet either of the following four criteria: (1) Annotated as disease-causing mutations (“DM”) by HGMD (50); (2) Annotated as pathogenic (“P”) or likely pathogenic (“LP”) with star 1+ (i.e., one submitter with assertion criteria, multiple submitters with assertion criteria, expert panel, or practice guideline) by ClinVar (51). Note, that the term “likely pathogenic” is used to mean greater than 90% certainty of a variant being disease-causing (84); (3) Annotated in DoCM (52); (4) Annotated in Catalog of Validated Oncogenic Mutations by CGI (53). We curated the inheritance patterns of CRMs and the tissues affected by lesions in disease.

We considered the following criteria to define a broader set of potentially clinically relevant gene sets: (1) HGMD genes (50): genes with at least one disease-causing (“DM”) variant, (2) ClinVar genes (51): genes with at least one variant denoted as pathogenic (“P”) or likely pathogenic (“LP”) with star 1+ by ClinVar. (3) DoCM genes (52): genes with at least one DoCM variant. (4) CGI genes (53): genes with at least one CGI variant. (5) OnkoKB genes (64): OncoKB™ Cancer Gene List includes both tumor suppressor genes and oncogenes. (6) CiVIC genes (65). (7) ClinGen haploinsufficiency genes (66): Dosage Sensitivity Curated Gene List was downloaded from ClinGen. Haploinsufficiency genes are defined as those with Haploinsufficiency Score of 3. (8) OMIM dominant and recessive, essential in culture, essential in mice: such gene lists were obtained from study (67).

#### Simulating mutations for distributions of ratio of missense to synonymous mutation

In our study of the distributions of expected ratio of missense to synonymous somatic mutations, we explored five distinct mutational models. These models were designed to simulate genetic mutations in genes where somatic mutations were observed, under the assumption that each coding location within a gene is equally likely to mutate.

Acknowledging that mutations frequently occur within trinucleotide contexts, we integrated features that conform to the patterns of observed somatic mutations, such as the mutational spectrum. In total, we considered five different mutational models:

1. Random Context Model (“Context-blind random model”): This model posits that each location in a gene has an equal likelihood of mutating to any other random variant allele, without the necessity for a specific trinucleotide context or nucleotide change.
2. Nucleotide Change Context Model (“REF>ALT context random model”): This model only requires the same nucleotide changes as observed in our cohort’s somatic mutations, regardless of the original trinucleotide context. For instance, if an ‘A’ to ‘G’ mutation is observed, this model would generate one ‘A’ to ‘G’ mutation without considering flanking bases.
3. Exact Trinucleotide Context Model (“BEFAFT context random model”): This model mandates the exact same trinucleotide context for a mutation to occur. For example, if a mutation is observed in the ‘AGT’ context, this model would only generate one random mutation (to either ‘A’, ‘T’, or ‘C’) in the same ‘AGT’ context.
4. Flanking Base Context Model (“BEF_AFT context random model”): This model solely matches the flanking bases of the trinucleotide context. For instance, if the observed mutation context is ‘AGT’, random mutations (to either ‘A’, ‘T’, or ‘C’) could occur in any context where ‘A’ and ‘T’ are the flanking bases (e.g., ‘ACT’, ‘ATT’).
5. Combined Nucleotide Change and Trinucleotide Context Model (“BEF[REF>ALT]AFT context random model”): This model is a fusion of the second and third models. It necessitates both the same trinucleotide context and the same nucleotide changes.

#### Detection of mutational signatures across tissues

We applied SignatureAnalyzer (40) to extract mutational signatures operative in normal tissues. This tool leverages non-negative matrix factorization and expectation- maximization algorithms. Specifically, SignatureAnalyzer was executed with the COSMIC reference signatures (v3.3.1_SBS_GRCh37) (9) as priors. The following parameters were used for the SignatureAnalyzer runs: --random-seed 0, -t spectra, --objective poisson, -- prior_on_H L1, --prior_on_W L1, -n 50, --max_iter 30000, --reference cosmic3.

Our initial approach involved aggregating all mutations into an “All Tissues” category. We then refined our focus to a ‘tissue’ level, categorizing samples from various donors into specific tissue groups and only including tissues with a minimum of 100 detected somatic mutations. We retained only those signatures with a minimum cosine similarity of 0.8. As a result, we extracted a total of 33 signatures across 24 tissues (Table S2).

#### Identification of multi-tissue mutations shared within the same donor

We defined shared somatic mutations as those observed across more than one tissue within a single donor. In contrast, unique somatic variants occur exclusively in a single tissue. The germ layer assignments for each tissue’s origin were obtained from Table S12 of study (32).

#### Patterns of shared mutations in two existing GTEx RNA-seq data sets

We reanalyzed two additional datasets from GTEx (30,32). Calls of somatic mutations from GTEx bulk RNA-seq data were taken from Table S3 of (30) and Table S7 of (32). Shared and unique somatic mutations were determined according to the definitions above.

#### Quantification and Statistical Analysis

##### Local distribution of somatic mutations across exomes

To examine the distribution of somatic variants across exomes, we divided CDS of each gene into non-overlapping 50-base pair windows. This fine-grained analysis was conducted using the Poisson Test for each gene, where the lambda parameter is derived from the ratio of the number of somatic mutations within the CDS to its length. Notably, the resulting *P*-values of each 50-bp window encapsulate the likelihood of observing an equal or greater number of somatic variants within a given window. Benjamini-Hochberg Procedure is utilized for multiple testing corrections. This method enables the identification of statistically significant deviations from the expected uniform distribution.

To visually articulate the comprehensive insights gained from this investigation, Manhattan plots are applied, providing an intuitive representation of the overall distribution and varaint clusters across the exomic landscape.

#### Somatic mutation burden and normalized burden

We defined the mutation burden of a sample as the count of somatic variants detected.

The normalized mutation burden of a sample was characterized as the mutation burden adjusted by the size of the sample’s exome regions (the number of bases with a minimum of 10x and a maximum of 5 times the median coverage).

#### Proportions of variance attributable to donors or/and tissues through ANOVA

We conducted an ANOVA to investigate the impact of donors and tissues on the normalized burden of somatic mutations per sample. By considering both donors and tissues as independent variables and integrating technical factors into the model, we aimed to account for potential confounding factors and gain an understanding of how donors and tissues contribute to the variance of normalized burden of somatic mutations. The regression model is shown in Formula 5.

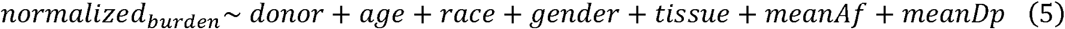

In our continued effort to estimate the variation attributable to donors, we drew comparisons with the approach of Rockweiler et al. (32) (detailed codes: https://github.com/conradlab/RockweilerEtAl/blob/main/figures/fig1.pzm_burden/input_files/get_rpt_estimates.R). Specifically, we employed a mixed-effects model, treating ‘donor’ as a random effect, as outlined in Formula 6. We used the R package rptR to fit the model and calculate the variation explained by donors, fitting Formula 6 using restricted maximum likelihood (77). For Gaussian data, we set the number of parametric bootstraps (nboot) to 500.

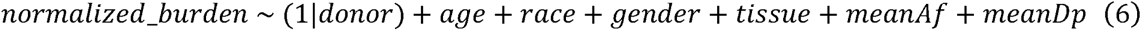

#### Evaluating the load of germline variants in CDS regions for individual genes

VCF of gnomAD variants was filtered to only include passing quality SNVs using BCFtools (74): ‘bcftools view -i ’FILTER=“PASS” && allele_type=“snv“’’. Two different allele frequency thresholds were used, requiring either alleles to be more frequent than 1% with bcftools view flags ‘--min-af 0.01’ or less frequent than 1% with ‘--max-af 0.01’.

Then a python script was used to (1) filter to retain only the longest transcript CDS per Ensembl gene ID and calculating the length from the gencode gtf file and (2) merge in the number of filtered gnomAD variants from the previous step using the Ensembl gene identifier and finally (3) calculate the Pearson correlation coefficient between CDS length and number of mutations for SNVs with either more or less than 1% frequency.

#### Utilizing both linear mixed models and linear regression to analyze and model the proportions of CpG mutations

We employed both a linear mixed model (LMM, Formula 7) and standard linear model (LM, Formula 7) to model the correlation between the proportions of CpG>T transitions and age. The *P*-values were derived from applying the “drop1()” function to the model with the test parameter set to “Chisq”. The models were structured as follows:

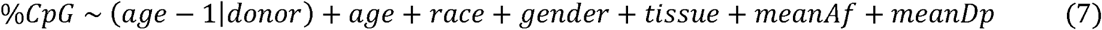

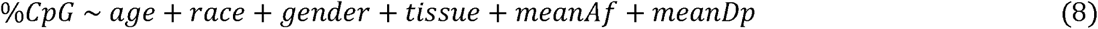

To more effectively investigate the interplay between CpG>T transitions, age, and cell proliferation, we categorized tissues with the highest 30% of cell proliferation scores as proliferative, with the remainder classified as non-proliferative. We fitted both LMM (Formula 9) and LM (Formula 10) independently within proliferative and non-proliferative tissues. The models were defined as follows:

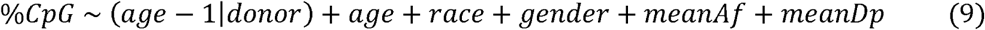

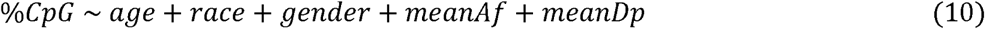

To ensure our conclusions are not biased by the cell proliferation cutoff, we adjusted the cutoff across 9 tertiles. We also modeled proportions of non-CpG C>T transitions as the response variable for contrast.

#### Enrichment of somatic mutations in the DHS, CTCF and pELS region

To test whether detected somatic mutations are enriched to occur in known cis-regulatory regions such as DHS, CTCF binding regions, or proximal enhancers, ENCODE SCREEN data for GRCh38 was downloaded on September 6th 2023 (68). This dataset contains genomic positions of numerous types of candidate cis-regulatory sequences (cCREs). Using LiftOver (76), we converted the genomic coordinates to the GRCh37 reference. Analysis was restricted to DHS regions (which comprise the entire dataset), and its subsets, which include CTCF binding regions and pELS that occur within 2kb of a transcription start site. If a single genomic region had multiple annotations, it was included in each of the analyses.

Since DHS regions correspond to open euchromatin, one hypothesis is that somatic mutations would be enriched to occur in these regions. We tested this by building a two-by-two contingency table with the number of sufficiently-covered loci that were or were not called variants and were or were not annotated as DHS binding sites, followed by a Chi-square test. Sufficiently-covered loci were those with enough depth to potentially call somatic mutations, which varied for each sample. This test resulted in a significant Chi-square *P*- value of 3.4×10^-5^, indicating a higher enrichment for somatic mutations in DHS regions. An additional bootstrapping test, which does not rely on the assumptions required for the Chi-square test. This involved randomly redistributing somatic mutation calls across sufficiently-covered loci and then calculating the number of somatic SNVs landing in DHS regions. This process was repeated for 7,000 iterations to generate a null-distribution to compare against the empirical counts. The bootstrapped *P*-value was 0 as not a single iteration had greater than or equal to the empirically observed 1,110 somatic mutations in DHS regions.

Performing the same tests as described above for CTCF regions, which resulted in a *P*-value of 2.0×10^-4^ for the Chi-square test and a *P*-value of 0 for the bootstrapping test.

For pELS, a class of cCREs, somatic variants were not enriched in annotated regions, with a Chi-square *P*-value of 0.53 and a bootstrap *P*-value of 0.78.

#### Associating CRMs with gene expression

We investigated the relationship between CRMs and gene expression levels using RNA-seq data. Median gene-level TPM by tissue was downloaded from the GTEx Portal. To categorize genes based on expression levels, we applied cutoffs of 1and 10, resulting in three groups: low-expressed, intermediate-expressed, and high-expressed genes.

## Acknowledgments

We thank Selina Vattathil, Lu Chen, Michelle Chan, and John Storey for their valuable contributions and insightful discussions during this research. The authors are pleased to acknowledge that the work reported on in this paper was substantially performed using the Princeton Research Computing resources at Princeton University which is a consortium of groups led by the Princeton Institute for Computational Science and Engineering (PICSciE) and Office of Information Technology’s Research Computing. This work was supported in part by NIH Grants U01HG007591 and R01GM110068 to JMA.

## Author Contributions

J.M.A. and H.X. conceptualized and designed the project. J.M.A. oversaw the overall project.

D.A. handled sample processing and sequencing. H.X. performed the analysis and wrote the mSOMA software. R.B. and T.C. performed data preprocessing, code cleaning, and data management. R.B. created a Docker version of mSOMA and conducted the analysis for Fig. 5C-E. C.M. contributed to part of Fig. 3F. H.X. and J.M.A. handled manuscript writing, review, and editing, with input and feedback from all authors.

## Declaration of Interests

All authors declare no competing interests.

## Notes

### Competing Interest Statement

The authors have declared no competing interest.

