## Supplementary Materials for "Landscape of human protein-coding somatic mutations across tissues and individuals"

**Table S1**. **Information of the 14 GTEx donors studied.**

| Donor | Age | Sex | Num Tissues Sequenced | Total Coverage | Num SMs Discovered |
| --- | --- | --- | --- | --- | --- |
| PT-WHSE | 20 | Male | 10 | 1488 | 688 |
| PT-XQ3S | 26 | Male | 11 | 1454 | 526 |
| PT-XUZC | 31 | Female | 12 | 1493 | 517 |
| PT-1399R | 38 | Male | 15 | 2217 | 390 |
| PT-13FTW | 44 | Male | 15 | 2054 | 402 |
| PT-XV7Q | 49 | Female | 26 | 3453 | 1151 |
| PT-13D11 | 58 | Female | 19 | 2543 | 680 |
| PT-1211K | 63 | Female | 16 | 2373 | 380 |
| PT-12WSD | 64 | Female | 25 | 3678 | 391 |
| PT-XUJ4 | 64 | Female | 22 | 2676 | 809 |
| PT-11DXX | 66 | Female | 21 | 2908 | 665 |
| PT-11GSP | 66 | Female | 28 | 4260 | 696 |
| PT-13OW8 | 66 | Male | 29 | 4236 | 560 |
| PT-X4XX | 69 | Male | 16 | 2254 | 615 |

**Table S2. Mutational signatures detected in our data.**

| Signature | Tissue | Cosine Similarity |
| --- | --- | --- |
| SBS-6 | All Tissues | 0.83 |
| SBS-6 | Aorta | 0.82 |
| SBS-6 | Esophagus-Muscularis | 0.82 |
| SBS-6 | Lung | 0.8 |
| SBS-6 | Skin-Sun Exposed Lower Leg | 0.83 |
| SBS-7B | Skin-Sun Exposed Lower Leg | 0.91 |
| SBS-15 | All Tissues | 0.93 |
| SBS-15 | Adrenal Gland | 0.82 |
| SBS-15 | Aorta | 0.95 |
| SBS-15 | Hippocampus | 0.82 |
| SBS-15 | Mammary Tissue | 0.86 |
| SBS-15 | Colon-Sigmoid | 0.89 |
| SBS-15 | Colon-Transverse | 0.81 |
| SBS-15 | Esophagus-Gastroesophageal Junction | 0.89 |
| SBS-15 | Esophagus-Muscularis | 0.92 |
| SBS-15 | Heart-Atrial Appendage | 0.89 |
| SBS-15 | Heart-Left Ventricle | 0.86 |
| SBS-15 | Liver | 0.87 |
| SBS-15 | Lung | 0.92 |
| SBS-15 | Nerve-Tibial | 0.82 |
| SBS-15 | Ovary | 0.83 |
| SBS-15 | Pancreas | 0.83 |
| SBS-15 | Skin-Not Sun Exposed Suprapubic | 0.85 |
| SBS-15 | Skin-Sun Exposed Lower Leg | 0.88 |
| SBS-15 | Small Intestine Terminal Ileum | 0.85 |
| SBS-15 | Spleen | 0.89 |
| SBS-15 | Stomach | 0.84 |
| SBS-15 | Testis | 0.88 |
| SBS-15 | Thyroid | 0.94 |
| SBS-15 | Vagina | 0.9 |
| SBS-18 | All Tissues | 0.91 |
| SBS-29 | All Tissues | 0.88 |
| SBS-95 | All Tissues | 0.89 |

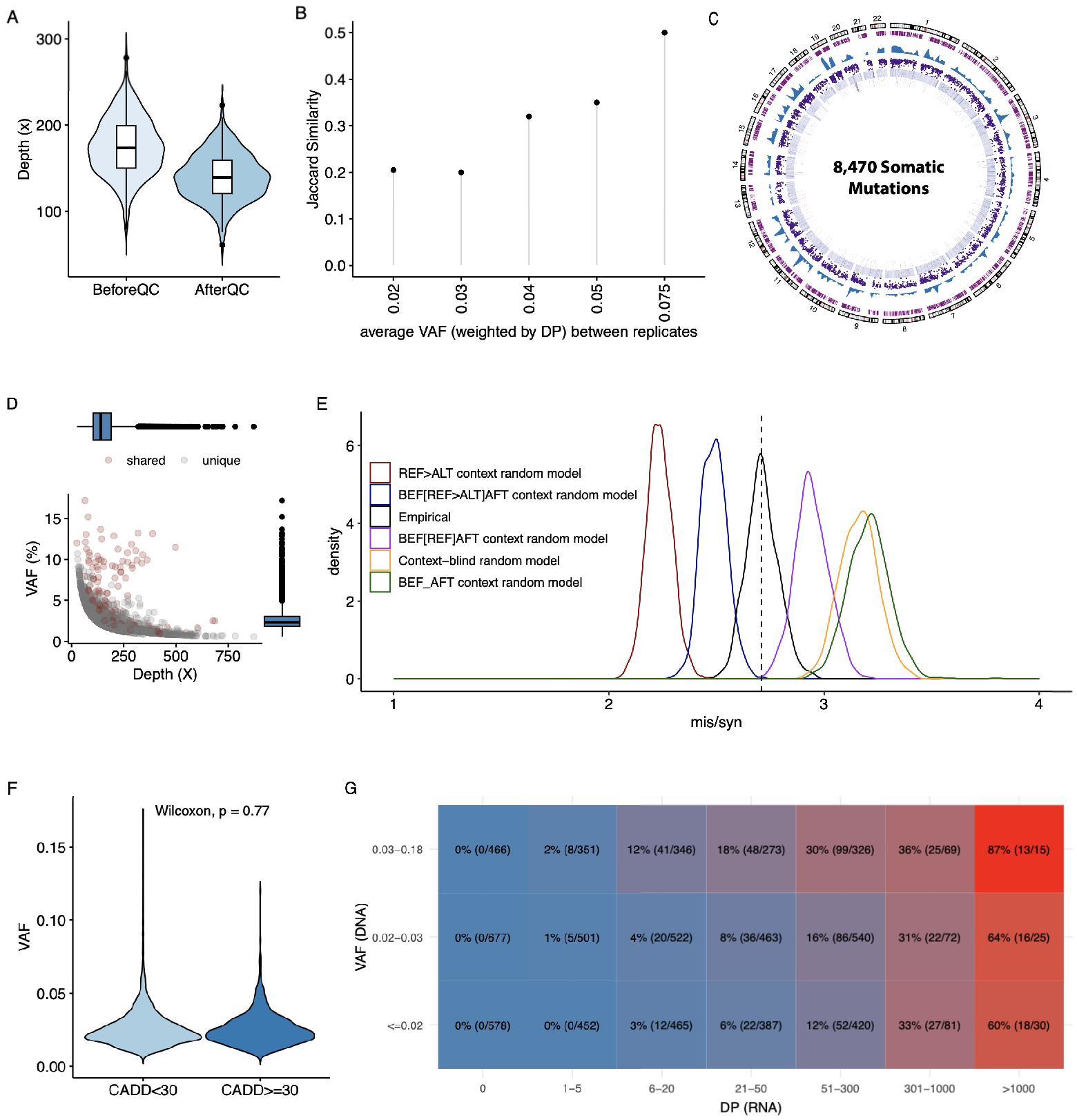

Fig. S1. Detection and validation of somatic mutations in our study.

(A) The average sequencing coverage before and after quality control for a total of 265 samples.
(B) Correlation between the average Variant Allele Frequency (VAF) and the Jaccard Similarity among replicate tissue samples. The average VAFs are computed as VAFs weighted by Depth (DP) across these replicate samples. Each data point in the plot represents a unique average VAF along with its corresponding Jaccard Similarity. The observed positive correlation suggests that an increase in the average VAF is associated with an increase in the Jaccard Similarity. This implies that higher VAFs have a higher likelihood of consistent detection across replicate samples.
(C) A Circos plot, created using the R Circlize package (https://doi.org/10.1093/bioinformatics/btu393), providing a panoramic view of the somatic mutations identified from WES samples. The outermost track depicts the chromosome ideogram with centromeres marked by red dashes. The next track depicts the callable region in purple which confines where somatic mutations could be detected. The blue density plot shows the concentration of discovered somatic mutations throughout the genome with added granularity from the purple "rainfall" plot which shows the location of individual mutations. The second-most and innermost two tracks represent the counts of reference alleles and alternate alleles as heatmaps. Loci with VAF < 0.05 are masked.
(D) Relationship between VAFs and Sequencing Coverage. Dot color denotes whether a variant is distinct or shared in more than one tissue of the same donor.
(E) Distributions of empirical and expected ratios of missense to synonymous somatic mutations under five context-specific models. Observed ratio in our data is indicated by a dashed black line.
(F) Comparison of distributions of VAF between deleterious (CADD>=30) and non-deleterious (CADD<30) variants.
(G) Changes in validation rates of somatic mutations identified in DNA-seq samples via RNA-seq, depicted in terms of VAFs in the DNA-seq (y-axis) and sequencing coverage in RNA-seq data (x-axis).

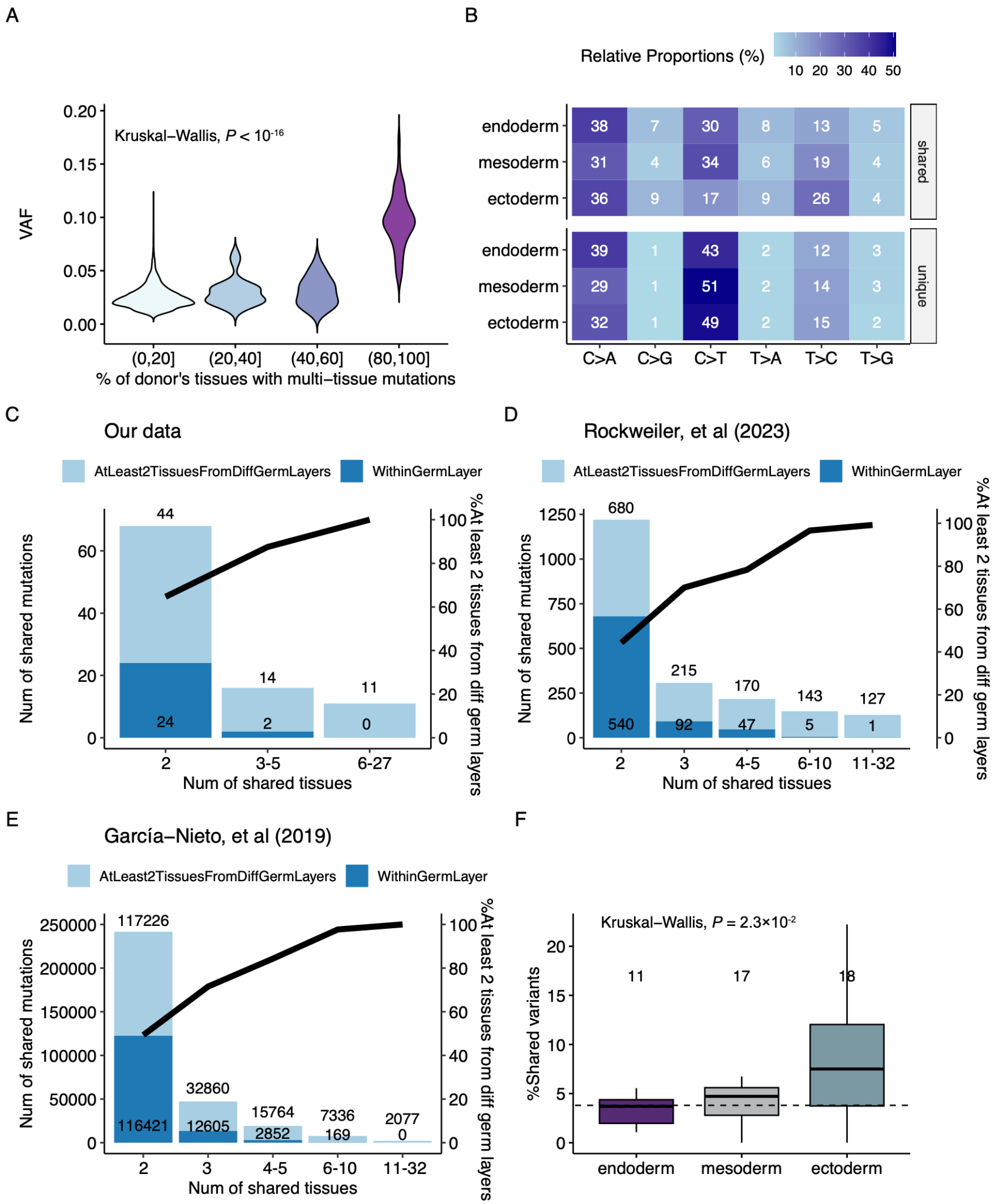

Fig S2. Patterns of shared multi-tissue mutations from the same donor.

(A) Relationship between VAF and the proportion of the donor’s tissues exhibiting the multi-tissue mutation.

(B) Mutational spectrum across germ layers, distinguishing between shared (top) and unique (bottom) mutations, with cell color denoting relative proportions.

(C-E) The left y-axis represents the numbers of multi-tissue mutations present in two or more tissues within the same germ layer (colored in dark blue) or in at least two tissues from different germ layers (colored in light blue). The right y-axis shows the proportions of multi-tissue mutations found in at least two tissues originating from different germ layers (light blue). The x-axis shows the number of tissues with shared mutations. Data were drawn from our study (C), Rockweiler et al. (2023) (D), and García-Nieto et al. (2019) (E).

(F) Comparisons of proportions of shared mutations per tissue across three germ layers. The aggregate number of sequenced tissues per germ layer is indicated at the top. A dashed line represents the overall proportion among 265 samples.

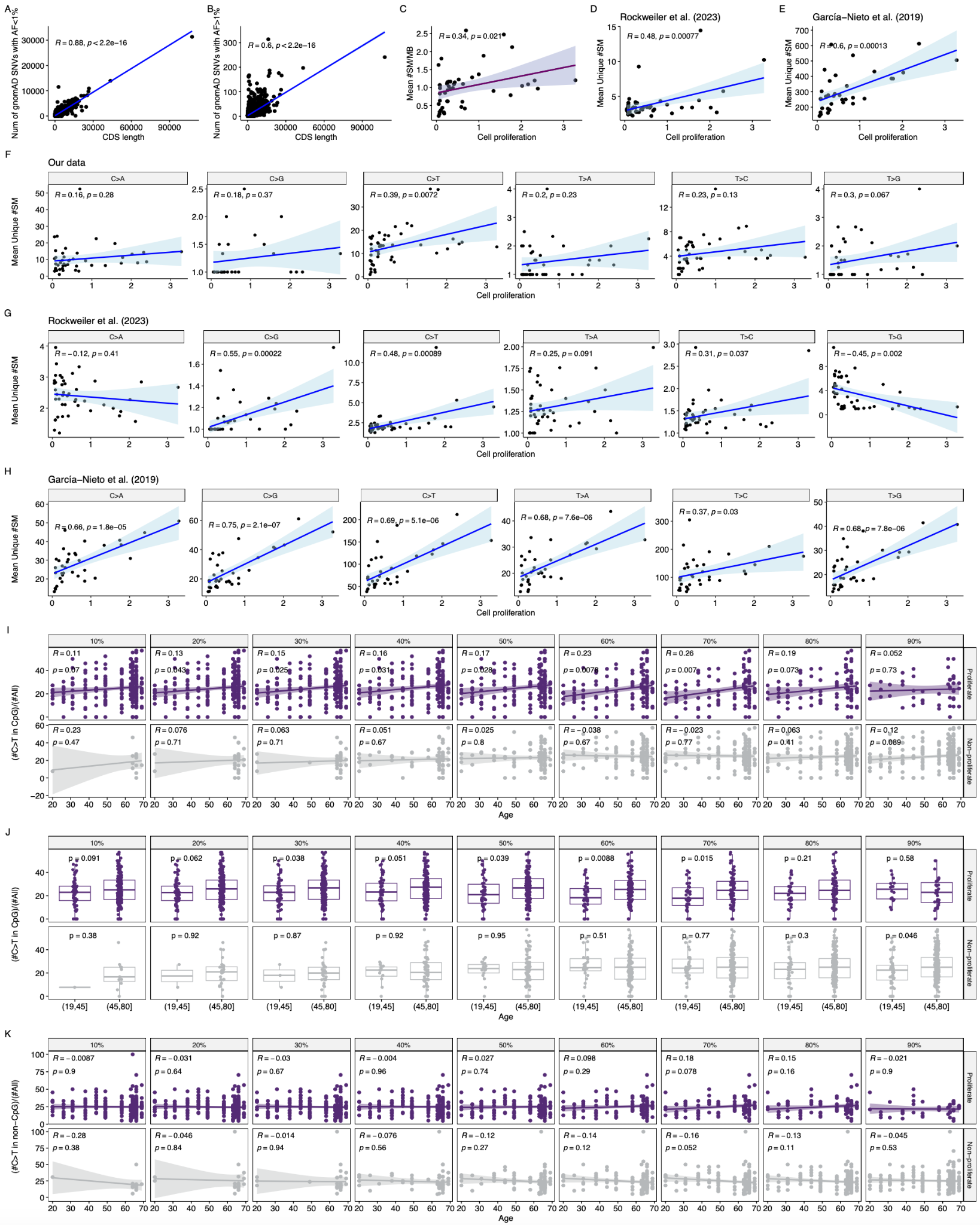

Fig S3. Factors that influence somatic mutations burden. Unless explicitly stated, the Pearson *R* and P-value are computed from linear regression and are denoted accordingly.

(A-B) Association between the length of the coding DNA sequence (CDS) and the gene-level impact of gnomAD variants with VAFs falling beneath 1% (A) and exceeding 1% (B).

(C) Relationship between tissue-specific cell proliferation and the average normalized overall burden across a variety of tissues

(D-E) Correlations between tissue-specific cell proliferation and the average number of unique somatic mutations across various tissues, as reported in the studies by Rockweiler et al. (2023) (D) and García-Nieto et al. (2019) (E).

(F-H) Association between tissue-specific cell proliferation and the average tally of unique somatic mutations (disaggregated by nucleotide changes) across diverse tissues, as observed in our data (F), and as reported in the studies by Rockweiler et al. (2023) (G) and García-Nieto et al. (2019) (H).

(I-J) Relationship between age and the proportions of CpG>T transitions between proliferative and non-proliferative samples. The facets correspond to different thresholds employed to differentiate between proliferative and non-proliferative samples. Age was treated as a continuous variable (I) or a categorical variable (J). The P-values in (J) were computed employing the Wilcoxon Test.

(K) Correlations between age and the proportions of C>T transitions that are not followed by G, between proliferate and non-proliferate samples.

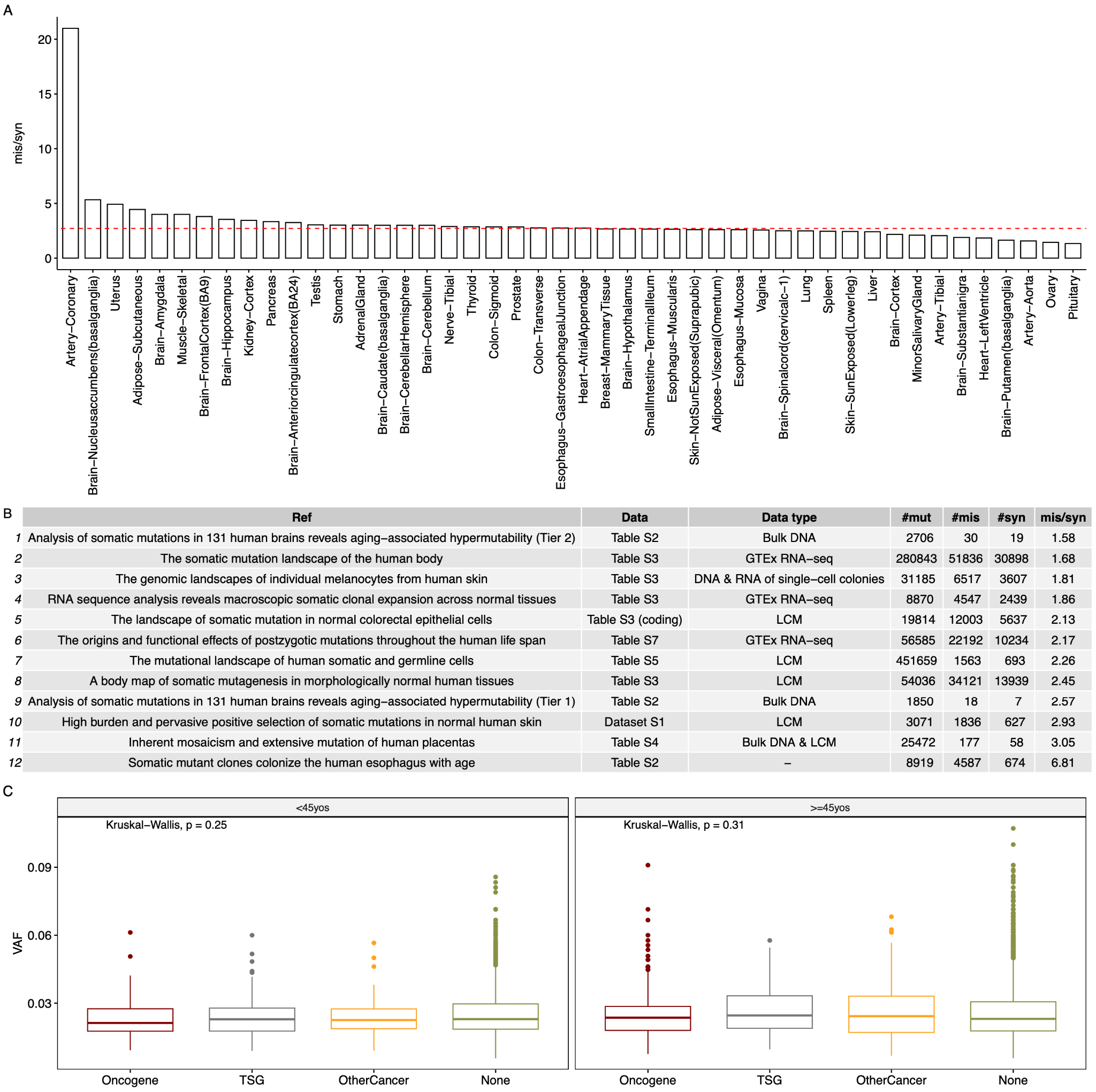

Fig. S4. Analysis of ratios of missense to synonymous variants and VAFs.

(A) Ratios of missense to synonymous variants across tissues. A red dashed line indicates the overall ratio of missense to synonymous variants across 265 samples.
(B) Ratios of missense to synonymous variants across Studies: The studies are ordered according to ascending ratios. The abbreviations used are as follows: #mut: Number of variants; #mis: Number of missense variants; #syn: Number of synonymous variants; mis/syn: Ratios of missense to synonymous variants.
(C) Distributions of VAFs for unique variants found in different categories of genes: oncogenes, tumor suppressor genes (TSG), other genes related to cancer (OtherCancer), and genes not associated with cancer (None).
